# Extensive hybridisation between multiple differently adapted species may aid persistence in a changing climate

**DOI:** 10.1101/2023.01.25.525480

**Authors:** I Satokangas, P Nouhaud, B Seifert, P Punttila, R Schultz, MM Jones, J. Sirén, H Helanterä, J Kulmuni

**Affiliations:** Organismal & Evolutionary Biology Research Programme, University of Helsinki, Viikinkaari 1, P.O.Box 65, 00014 University of Helsinki, Finland; Department of Entomology, Senckenberg Museum für Naturkunde, Am Museum 1, D-02826 Görlitz, Germany; Biodiversity Centre, Finnish Environment Institute, Helsinki, Finland; Institute of Biotechnology, HILIFE - Helsinki Institute for Life Science, 00014 University of Helsinki, Finland; Ecology and Genetics research unit, University of Oulu, P.O. Box 3000, 90014 University of Oulu, Finland; Tvärminne Zoological Station, University of Helsinki, J.A. Palménin tie 260, FI-10900 Hanko, Finland

**Author notes:** **Author for correspondence:** Jonna Kulmuni Ina Satokangas.

**Keywords:** Speciation, Hybridisation, Adaptation, Hymenoptera, *Formica* wood ants, Mosaic hybrid zone

## Abstract

Hybridisation and gene flow can have both deleterious and adaptive consequences for natural populations and species. To better understand the extent and consequences of hybridisation in nature, information on naturally hybridising non-model organisms is required, including characterising the structure and extent of natural hybrid zones. Here we study natural populations of five keystone mound-building wood ant (*Formica rufa* group) species across Finland. No genomic studies across the species group exist and the extent of hybridisation and genomic differentiation in sympatry is unknown. Combining genome-wide and morphological data, we show that *Formica rufa*, *F. aquilonia*, *F. lugubris*, and *F. pratensis* form distinct gene pools in Finland. We demonstrate more extensive hybridisation than previously thought between all five species and reveal a mosaic hybrid zone between *F. aquilonia*, *F. rufa* and *F. polyctena*. We show that hybrids between these climatically differently adapted species occupy warmer habitats than the cold-adapted parent *F. aquilonia*. This suggests hybrids occupy a different microclimatic niche compared to the locally abundant parent. We propose that wood ant hybridisation may increase with a warming climate, and warm winters, in particular, may provide a competitive advantage for the hybrids over *F. aquilonia* in the future. In summary, our results demonstrate how extensive hybridisation may help persistence in a changing climate. Additionally, they provide an example on how mosaic hybrid zones can have significant ecological and evolutionary consequences because of their large extent and independent hybrid populations that face both ecological and intrinsic selection pressures.

## Introduction

Hybridisation and gene flow between closely related species are common in nature and predicted to increase because of human activities and warming climate (Chunco, 2014; Mallet, 2005; Rieseberg, 2009; Scheffers et al., 2016). Changing environmental conditions induce range shifts and connections between previously allopatric populations, leading to increased rates of hybridization that may over longer evolutionary time scales either collapse species barriers (Owens & Samuk, 2019) or reinforce them (Lemmon & Juenger, 2017; Pfennig, 2016).

In terms of fitness, hybridisation can result in both positive and negative effects. New genetic variation acquired from introgression has been repeatedly shown to underlie adaptation into new niches (De-Kayne et al., 2022; Kagawa & Takimoto, 2018; Meier et al., 2017), and facilitate range shifts (Leroy et al., 2020), but hybridisation can also result in deleterious fitness effects (Ålund et al., 2013) and mortality (Ellison et al., 2008). Revealing the extent and consequences of hybridisation in natural populations provides essential knowledge on the fate and adaptive potential of hybrid populations. Thus, it is important to investigate how hybridisation in natural populations may change with warming climate.

The impact of hybridisation is further influenced by its geographical extent, and hence the type of the hybrid zone: In contrast to more commonly studied and geographically restricted clinal zones (Barton & Hewitt, 1985), mosaic hybrid zones may be as large as the species’ sympatric area. In mosaic zones (Arnold, 1997), hybridisation events are typically independent of each other and admixed populations alternate in space with non-admixed and backcrossed populations (Arnold, 1997; Harrison, 1986; Rand & Harrison, 1989). In the mosaic zone model, the parental species are assumed to be adapted to different environments, and hybrids may either have low fitness (due to intrinsic selection), or be favoured (extrinsic, environment-dependent selection) (Arnold, 1997). Mosaic hybrid zones have been reported, e.g., for different plant species (Abbott & Brennan, 2014; Rieseberg et al., 1999) and molluscs (Fraïsse et al., 2014). Since mosaic zones can be large, and selection can act independently in different hybrid populations, they may result in more heterogeneous patterns and geographically more widespread effects in contrast to clinal hybrid zones. Furthermore, independently evolving hybrid populations allow studying predictability in the outcomes of hybridisation (Nouhaud et al., 2022).

Hybridisation is commonly investigated between two species. However, multi-species hybridisation has been documented from white oaks (Reutimann et al., 2020) to birds (Grant & Grant, 2020; Natola et al., 2022; Ottenburghs, 2019), fish (Banerjee et al., 2022), and butterflies (Heliconius Genome Consortium, 2012). Multi-species hybridisation is likely to be more frequent in sympatric groups with a low degree of reproductive isolation, and one species may act as a “conduit” species and a vehicle of gene flow between two otherwise isolated species (Grant & Grant, 2020).

Wood ants of the *Formica rufa* species group (Hymenoptera, Formicidae) represent ideal study organisms to understand hybridisation in nature: they have diverged recently, within the past 500.000 years (Goropashnaya et al., 2004, 2012; Portinha et al., 2022) and have a wide sympatric range in Eurasia. Indeed, several species hybridise extensively within the sympatric range based on morphological data (Seifert, 2021). Furthermore both pre- and post-zygotic isolation, including intrinsic and environmentally dependent selection, have been detected within the group (Kulmuni et al., 2010; Kulmuni & Pamilo, 2014; Seifert, 2018). Specifically, hybridisation has resulted in male-biased mortality, female heterosis and potential for thermal adaptation (Kulmuni & Pamilo, 2014; Martin-Roy et al., 2021). Understanding hybridisation in the *F. rufa* group ants is important, since they are keystone species in boreal and mountain forests across Eurasia (Stockan & Robinson, 2016; Trigos-Peral et al., 2021). They build stable nest mounds (Seifert, 2018), and polygynous (i.e., multiple queens per nest) species and populations in particular can reach densities of hundreds of nests per km², as well as impact ecosystem characteristics, ranging from nutrient cycling below ground to aphid farms in the forest canopy. In Finland, at least five species of the *F. rufa* group have been reported to occur in sympatry and to be ecologically similar enough for potential hybridisation: *Formica rufa* (Linnaeus, 1761), *F. aquilonia* (Yarrow, 1955), *F. polyctena* (Förster, 1850), *F. lugubris* (Zetterstedt, 1838), and *F. pratensis* (Retzius, 1783). Of these, multiple hybrid populations of *F. aquilonia* and *F. polyctena* have been characterised genetically within a small region in southern Finland (Beresford et al., 2017). In addition, morphological and allozyme data suggests several other hybrid findings between *F. aquilonia* and *F. polyctena,* mainly in southern Finland (Pamilo & Kulmuni, 2022; Seifert, 2021; Sorvari, 2022), and indications of *F. aquilonia* and *F. lugubris* hybridisation in northern Finland (Pamilo & Kulmuni, 2022). However, outside of *F. aquilonia* and *F. polyctena* (Nouhaud et al., 2022; Portinha et al., 2022) no genomic studies on the *F. rufa* group exist, and consequently no information on the congruence between morphological and genomic data is available.

Under a changing climate, hybridisation between differentially adapted species may provide an opportunity to reshuffle adaptations and help species to cope with environmental shifts. The five *F. rufa* group species we focus on here differ in their life history characteristics and habitat preferences (Seifert, 2018), and based on their Eurasian distributions, are expected to be adapted to different climatic conditions (Table 1). We refer to northern *F. aquilonia* as cold-adapted and southern *F. polyctena* as warm-adapted, since in laboratory conditions *F. aquilonia* tolerates cold and *F. polyctena* survives in heat better than vice versa (Martin-Roy et al., 2021) Similar to *F. aquilonia*, *F. lugubris* has a boreo-alpine distribution and is likely cold-adapted. In contrast, *F. rufa* and *F. pratensis* live at lower altitudes, and have a more southern distribution, similar to *F. polyctena*, and are assumed to be warm-adapted (Table 1) (Seifert, 2018).

**Table 1.**
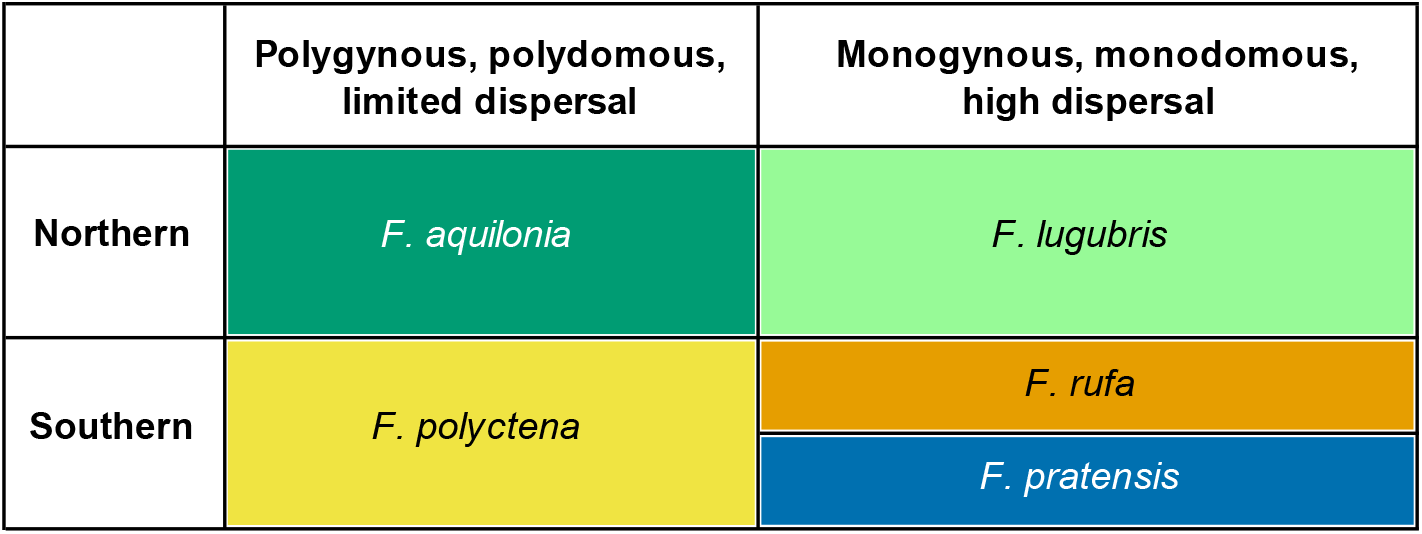
Characteristics of the study species in Finland.

Since successful wood ant hibernation, reproduction, and brood development are temperature-dependent, climatic conditions are critical for their fitness and survival through affecting within-nest temperatures throughout the year (Frouz, 2000; Frouz & Finer, 2007; Kadochová & Frouz, 2014). However, the ants can enhance their fitness by active nest thermoregulation from early spring until the autumn (Horstmann and Schmid 1986; Kadochová and Frouz 2014). The active regulation is paused for the hibernation period, but recovered in the spring, when the species start to reproduce at different but overlapping times (Seifert, 2018), with brood production and development continuing until summer. In the winter, too high hibernation temperatures increase ant metabolism, lead to reduced lipid reserves, and thus reduced fitness and increased mortality (Sorvari et al., 2011). Specifically, workers need the body fat reserves in early spring for nest heating (Seifert, 2018), and queens for successful sexual reproduction (Sorvari et al., 2011). Understanding the extent and consequences of hybridisation between wood ant species with different climatic and life history characteristics is important for understanding their survival. This is especially relevant near polar regions, like in Finland, where climate warming is relatively fast and leads to pronounced extreme weather phenomena (cold periods in the spring and warm winters). Specifically, hybridisation between warm- and cold-adapted species could provide a fitness advantage over the parental species when climate fluctuates. Indeed, an earlier laboratory study indicates that the natural hybrids of *F. aquilonia* and *F. polyctena* can tolerate warmer temperatures than *F. aquilonia* (Martin-Roy et al., 2021).

This study has three aims. First, we aim to detect whether sympatrically occurring *F. rufa* group wood ant species are genomically distinct, suggesting that barriers to gene flow exist and keep the gene pools separate even in sympatry. If morphologically distinguishable groups (i.e., the currently identified *F. rufa* group species) cannot be grouped based on genome-wide data, this would suggest the morphotypes are part of a larger panmictic gene pool. Second, we aim to characterise the extent of *F. polyctena* and *F. aquilonia* hybridisation across Finland, as well as any signs of hybridisation or introgression across the five species within the group. We expect this based on morphological signs of hybridisation across the *F. rufa* group across their sympatric Eurasian ranges (Seifert, 2018, 2021). Lastly, we assess whether specific environmental factors – temperature or precipitation that are crucial for the nest thermoregulation – correlate with the occurrences of *F. aquilonia* × *F. polyctena* hybrids. We hypothesise that these hybrids will occur in warmer and drier habitats than non-admixed cold-adapted *F. aquilonia* in Finland. This would support the hypothesis that hybrids may better cope with a warming climate. Combined with information on observed and expected heterozygosity, we discuss the potential for range expansions and adaptation to environmental changes by wood ant hybrids.

## Material & Methods

### Sample collection

To genomically characterise the five sympatric wood ant species and the extent of hybridisation and introgression among them, we used samples collected throughout Finland. These were complemented by already published samples from Scotland and Switzerland (Portinha et al., 2022), to provide a broader view.

In total, 68 of the nests are located in Finland, and one in Russia, each belonging to a separate population. We also included 20 previously published genomes of *F. aquilonia*, *F. polyctena* and admixed individuals sampled from Finland (eight nests and populations), Switzerland (three *F. aquilonia* nests from two populations and 6 *F. polyctena* nests from four populations), and Scotland (three *F. aquilonia* nests from two populations) (Portinha et al., 2022).

Altogether the ant data consists of samples from 89 nests of five *F. rufa* group species: *F. aquilonia*, *F. polyctena*, *F. rufa*, *F. lugubris* and *F. pratensis*. For the genomic analyses, a single ant per nest (and population for Finnish nests) was used. The morphological analysis was carried out on all newly sampled Finnish and Russian nests. Morphological data was available also for some of the previously published genomes (Table 2).

**Table 2.** Number of individuals with genomic data used in this study. NUMOBAT data available = “NUM”

|  | Finnish samples | Swiss samples | Scottish samples | Russian samples |
| --- | --- | --- | --- | --- |
| Genomes from Portinha et al. 2022 | 8 (NUM= 4) | 9 (NUM =2) | 3 (NUM=1) | - |
| Samples analysed for this study | 68 (NUM: 68) | - | - | 1 (NUM=1) |

The majority of the Finnish samples originate from the National Forest Inventory (NFI years 2005-2008; (Punttila & Kilpeläinen, 2009)), supplemented by samples from individual researchers. Samples were chosen based on initial morphological evaluation aiming at eight or more samples (and hence populations) per species. However, more detailed morphological investigation (NUMOBAT, Numeric Morphology-Based Alpha-Taxonomy, see below) was carried out after choosing the samples. Moreover, we aimed at covering the known distribution of each species in Finland (Stockan & Robinson, 2016), and the samples were chosen to be as evenly distributed across each species’ distribution in Finland as possible. To better characterise the extent of hybridisation between *F. polyctena* and *F. aquilonia* across Finland, we included more samples from these species (and putative admixed samples).

### Morphological data

For the morphological species identification multiple individuals from each nest were analysed using NUMOBAT, which produces species assignment probabilities at the nest level. Altogether 17 morphological characteristics were measured for three to six individual ants per nest (See Suppl Table 1 for a description of the characteristics). The recording of NUMOBAT characters is described in detail in Seifert (2021).

All 76 samples for which NUMOBAT data was available (69 nests collected for this study and seven nests analysed in (Portinha et al., 2022), see Table 2), were run in a Linear Discriminant Analysis (LDA). All 73 Finnish and Russian samples were run as wild-cards (i.e., without imposing a species hypothesis), and three non-Finnish samples with previously allocated species hypotheses, together with a reference sample of 1996 workers of 406 Palaearctic nest samples with pre-established species hypotheses. The reference samples contained only samples of the five species dealt with here which did not show phenotypical indications of hybrid identity. There was only a small amount of reference data for *F. lugubris* and *F. pratensis* in six of the morphological characteristics. Hence, all samples were first run in a 5-class LDA using 11 morphological characteristics for which sufficient reference data was available (see also Suppl Table 1). This led to a fixation of posterior probabilities for samples assigned as *F. lugubris* or *F. pratensis* in this 1st step of analysis. In the second step, all samples that were not classified as *F. lugubris* or *F. pratensis* were run as wild-cards in a 3-class LDA with reference data considering all available 17 characters. The resulting posterior probabilities were then adjusted in a way that the sum of probabilities of all five species equals to 1.0.

The adjustment of all five probabilities to give a sum of 1.0000 was done as follows

a = P(*F. lugubris*)+P(*F. pratensis*) from the 1st step

b = 1-a

F. aquilonia, F. polyctena, and F. rufa: P(final) = b * P(2nd step)

F. lugubris, F. pratensis: P(final) = P(1st step)

To study the concordance of morphological and genomic species assignments, we computed per species means and standard deviations of the respective LDA values for all non-admixed individuals. For the individuals admixed between *F. aquilonia* and *F. polyctena*/ *F. rufa* clade, we performed Pearson’s correlation between the ADMIXTURE (K=5) and LDA values of *F. aquilonia*, and for the admixed *F. lugubris* individuals, the respective *F. lugubris* values. The lack of variance in ADMIXTURE values prevented testing this statistical correlation for the subset of non-admixed individuals used in ADMIXTURE.

### Genomic data

For the genomic characterisation of our study species and individual samples, as well as studying the extent of hybridisation using nuclear admixture and mito-nuclear mismatch, we performed whole-genome sequencing for our 69 new study samples and analysed these along with 20 samples previously sequenced (Portinha et al., 2022).

#### DNA extraction and sequencing

DNA was extracted with Qiagen DNeasy extraction kit using a single ant per nest and population, according to the manufacturer protocol for insects. DNA libraries were constructed and sequenced at the Biomedicum Functional Genomics Unit (FuGU), with NEBNext Ultra II FS DNA library preparation and Illumina NovaSeq S4: 2×150bp (1 lane) sequencing.

#### Variant identification

Raw Illumina reads and adapter sequences were trimmed using TRIMMOMATIC (v0.39; parameters LEADING:10, TRAILING:10, SLIDINGWINDOW:4:15, MINLEN:50; (Bolger et al., 2014)) before mapping against a Finnish *F. aquilonia × F. polyctena* hybrid reference genome (Nouhaud et al., 2021) using BWA MEM with default parameters (v0.7.17; (Li, 2013)). Duplicates were removed using PICARD TOOLS with default parameters (v2.21.4; http://broadinstitute.github.io/picard). Average read depth computed across individuals with MOSDEPTH (v0.3.3 (Pedersen & Quinlan, 2018)) after duplicate removal was 16.0× (Suppl Table 2).

##### Nuclear genomic data

Single nucleotide variants (SNPs) for the nuclear genome were called with FREEBAYES software (v1.3.1, -k option was used for disabling population priors; (Garrison & Marth, 2012)) and the VCF was normalised using VT (v2015.11.10; (Tan et al., 2015)). Sites located at less than two base pairs from indels were excluded, as well as sites supported by only Forward or Reverse reads, using BCFTOOLS (v1.10; (Danecek et al., 2021)). Multi-nucleotide variants were decomposed with the vcfallelicprimitives command from VCFLIB (v1.0.1 (Garrison et al., 2022)). The following steps were computed with BCFTOOLS. Only biallelic SNPs with quality equal or higher than 30 were kept. Sites were then filtered based on individual sequencing depth: individual genotypes that had higher than twice the mean depth of the individual in question were set as missing. Genotyping errors that would occur e.g. due to misaligned reads were removed by excluding sites that displayed heterozygote excess after pooling all samples (P < 0.01, --hardy command from VCFTOOLS v0.1.16, (Danecek et al., 2011)). Subsequently, individual genotypes with quality < 30 and depth of coverage < 8 were coded as missing data. Sites with more than 10% missing data over all samples were discarded. Finally, singletons were removed (minor allele count < 2) with VCFTOOLS (v0.1.16).

After these steps 1.829.565 SNPs across 89 individuals remained in the final dataset with sequencing depth of at least 8×. All the genetic analyses (detailed below) were performed subsampling 9770 genome-wide SNPs that were obtained by thinning the final dataset with 20kb distances to minimise linkage disequilibrium, as well as by excluding chromosome number three (social chromosome with reduced recombination due to supergene inversions (Brelsford et al., 2020)) and shorter contigs with no known genomic location.

##### Mitochondrial genomic data

SNP calling for the mitochondrial genome was done with FREEBAYES and a frequency-based approach (--pooled-continuous option). The SNP filtering followed the same pipeline that was used for the nuclear genome except the steps after keeping biallelic SNPs with quality equal or higher than 30 were not applied for the mitochondrial data. This resulted in 248 biallelic SNPs across the 89 individuals. Individual FASTA files were created using vcf-consensus (VCFTOOLS v0.1.16) and aligned with MAFFT (v7.505, (Katoh & Standley, 2013)).

We used two lines of evidence to study the genome-wide differentiation of the studied species and to identify admixed individuals and the extent of introgression: individual nuclear genomic ancestry proportions with ADMIXTURE, and mito-nuclear mismatches. To support these, we utilised phylogenetic network and principal component (PCA) analyses. All genomic analyses were used in combination with results from the NUMOBAT analysis that allowed morphological identification of the species.

#### Admixture analysis

ADMIXTURE (Alexander et al., 2009) was performed with the 9770 nuclear SNP data to study the species’ genomic distinction and admixture. All five species were included, but we aimed at balanced sample sizes of non-admixed individuals (from six to eight individuals per species) and thus randomly selected six *F. aquilonia* individuals, since unequal sample sizes could yield biased clustering results (Toyama et al., 2020). For ADMIXTURE, the number of clusters K was chosen to range from K=2 to K=5, with prior knowledge of sampling from five morphologically identified species. Random selection of six *F. aquilonia* individuals was repeated multiple times to confirm that results were not driven by certain individuals. The most likely value of K was chosen based on cross-validation error (CV, prediction of withheld data (Alexander & Lange, 2011)).

#### Mitotype network

To further study genetic structure and divergence, and to detect potential mito-nuclear mismatches we characterised the mitotypes of our samples using the mitochondrial SNP data. Samples of the same species are expected to have mitochondrial haplotypes that cluster together. Mito-nuclear mismatch, i.e., that the mitochondrial and nuclear genomes of an individual originate from different species, would be an indication of hybridisation and introgression. A mitochondrial median-joining network was created with POPART (Leigh & Bryant, 2015).

#### Phylogenetic network analysis

Using the nuclear genomic SNP data, we constructed a phylogenetic network to support other analyses of genomic distinctness and hybridisation. In the phylogenetic network, individuals that are assigned to the same species should form clusters despite their geographical location. If the samples do not form separate species-specific clusters but samples from different species cluster together, alternating with each other, it is indicative of a possible eco- or morphotype level difference instead of genomically distinct species. If no clear clusters exist at all, it indicates extensive gene flow between morphologically differentiated samples. If an individual is located in between species’ clusters, and does not thus cluster with other samples, it is a putative signal of hybridisation and introgression.

For the phylogenetic network, we computed pairwise genetic distance between all pairs of samples using the script distMat.py from github.com/simonhmartin/genomics_general. A phylogenetic network was computed from the distance matrix with SplitsTree (version 4.17.1, (Huson & Bryant, 2006)), the NeighbourNet approach (Bryant & Moulton, 2004) and default parameters.

#### PCA

For additional analysis of the genetic structure, genomic distinction, and hybridisation of the species, we performed a principal component analysis (PCA) using the nuclear genomic SNP data. Similarly to the phylogenetic network, genomically diverged species are recognised from forming separate clusters, in contrast to e.g. geography to drive the clustering patterns. PCA was performed with PLINK (v2.00a3, (Purcell et al., 2007)).

#### Inbreeding coefficients

We additionally studied the inbreeding coefficient of our Finnish samples to see if the admixed individuals harbour elevated heterozygosity in comparison to non-admixed individuals. Lower inbreeding coefficients and thus higher heterozygosity in hybrids may allow better adaptation to changing climatic conditions. Calculation of the per individual inbreeding coefficients was performed with VCFTOOLS (v0.1.16) --het command, for all Finnish and the one Russian sample (N=77) pooled together. We used a Wilcoxon Rank Sum test to test the difference in inbreeding coefficients between hybrid individuals and individuals of the parental species.

### Climate data

#### Ambient temperature and precipitation

To study whether temperature or precipitation correlate with the occurrence of the hybrids between *F. aquilonia* and *F. polyctena*, and specifically, if the hybrids occur in warmer and drier habitats than cold-adapted *F. aquilonia,* we analysed climate data for the Finnish nest locations. To restrict the comparison to the sympatric range, all nests north of the northernmost *F. rufa* locality (in the absence of non-admixed *F. polyctena* populations) at 63.36 degrees latitude, were excluded from this analysis, leaving a total of 33 nest sites (see Fig 1B). We generated climatic time series for each of these nests extending 1, 5, and 20 years backwards from the nest-specific field sampling date. Time series data for temperature were extracted from digital rasters of daily mean air temperature, and for precipitation from digital rasters of daily precipitation sum (sourced from the Finnish Meteorological Institute ClimGrid gridded daily climatology at a 10 km spatial resolution; (Aalto et al., 2016)) based on nest site coordinates.

**Figure 1.**
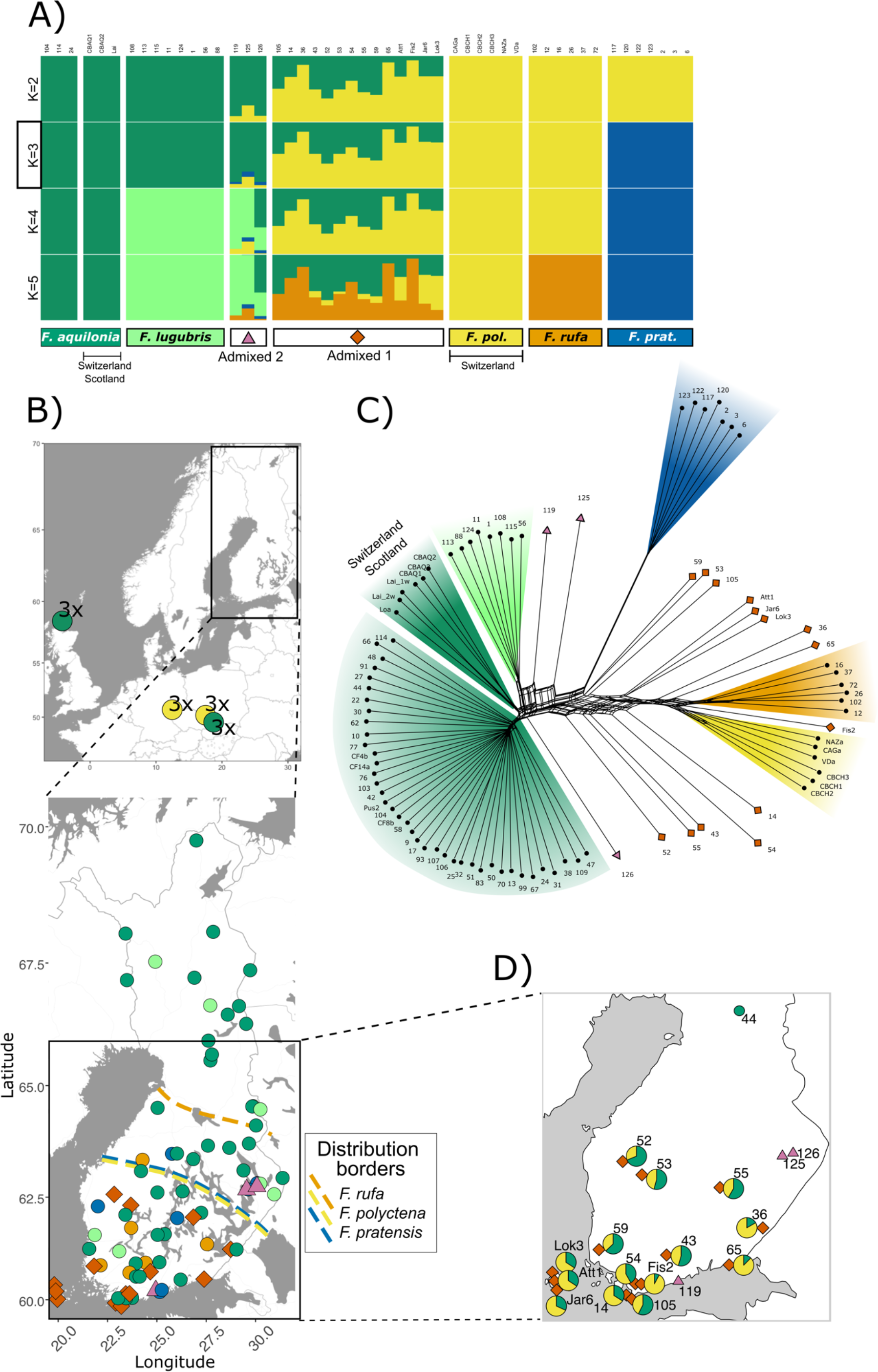
**A)** ADMIXTURE analysis recovers 5 species and admixture involving all species. Colored legend bars indicate non-admixed samples, while admixed individuals are divided into two groups based on their parental species. K=3 has the smallest cross-validation error. Depending on which *F. aquilonia* samples are chosen for the analysis, the order at which a distinction between sister species (*F. aquilonia* vs *F. lugubris*, and *F. polyctena* vs *F. rufa*) is seen within the clades varies, as do the *F. rufa* and *F. polyctena* contributions in hybrids. **B)** The upper map presents sampling locations from Scotland (3 × *F. aquilonia*), eastern Switzerland (3 × *F. aquilonia* and 3 × *F. polyctena*) and western Switzerland (3 × *F. polyctena*). The lower map shows sampling for the genomic and morphological data in Finland (one sample is located at the Russian side of Finland’s eastern border). Colours and symbols are as in panel A). The northern distributional borders are shown for species that do not inhabit the whole country, as in Stockan and Robinson (Stockan & Robinson, 2016). **C)** Phylogenetic network based on absolute genetic pairwise distances computed over nuclear SNPs supports the results of the ADMIXTURE analysis. Note sample 44, which is nuclearly non-admixed *F. aquilonia,* but has mito-nuclear mismatch. **D)** Sampling locations with individual IDs for all samples that show admixture or mito-nuclear discordance, and ADMIXTURE proportions (K=3) for the admixed samples of “Admixed 1” group, i.e., individuals admixed between *F. aquilonia* clade and *F. polyctena* clade. As in panel A), K=3, green indicates the *F. aquilonia/ F. lugubris* clade and yellow the *F. polyctena/ F. rufa* clade.

For each nest and year in the time series, we calculated annual winter (hibernation) and spring (reproduction) mean temperatures and precipitation from the daily values. The winter hibernation season was defined as the months of December to February and the spring reproductive season as April and May. To represent the warmest conditions experienced in winter at each site, we used the upper quantile of winter temperatures and the number of days when temperature reached over 0°C. To represent the coolest conditions experienced in spring we used the lower quartile of spring temperatures and the number of frost days. These values were used for each of the time periods in subsequent regression modelling.

To test whether *F. aquilonia* and hybrid nest sites diverged climatically within their sympatric range, we fitted Bayesian logistic regression models using the R package *brms* (Bürkner, 2017) with nest status (hybrid vs *F. aquilonia*) as the response variable. We fitted ten separate models with either one of the climatic covariates, or site latitude or longitude, as the explanatory variable. We did not include multiple explanatory variables in a single model due to high multicollinearity between the climatic variables and the coordinates, and the relatively low sample sizes. We normalised the explanatory variables to have mean zero and unit variance, and used an informative normal(0,2) prior distribution for the model parameters. The models were ranked according to their expected log pointwise predictive density from leave-one-out cross-validation using the *loo* R package (Vehtari et al., 2017). We also checked whether the exclusion of three nests that are spatial and climatic outliers, due to their location in the Åland islands in the Baltic Sea, altered the model results.

### Within-nest temperature

To study whether ambient temperature affects within-nest temperatures as experienced by the ants, we tested whether within-nest temperatures correlated with ambient temperatures or differed between hybrid and *F. aquilonia* nests. The populations used in these analyses were different from the ones used for the above morphological, genomic, and ambient climatic analyses.

In the years 2020 and 2021, we deployed Hobo data loggers to record within-nest temperatures hourly in 42 ant nests located within three admixed and two *F. aquilonia* populations in southern Finland (Suppl Fig 1). In total, four populations were sampled in 2020 and three populations in 2021; two of these were sampled in both years. In most cases, data logging ran continuously from 1st January until 31st July. However, data logging in one of the 2021 populations (‘Svanvik’) ran during May-July only. Daily mean within-nest temperatures per nest were calculated from the hourly values. Daily mean ambient temperatures for the corresponding dates were then extracted for each ant population based on the mean coordinates of nests within that population from the ClimGrid gridded daily climatology rasters at a 1 km spatial resolution (Aalto et al., 2016).

We furthermore modelled the mean daily within-nest temperatures of these nests in selected winter (January, February), spring (March-May) and summer (June, July) months as a function of nest status (hybrid vs *F. aquilonia*), site ambient temperature and sampling year (2020 vs 2021) using the R package *brms* (Bürkner, 2017). We compared a simple additive model structure (base model) with one including an interaction between nest status and ambient temperature (interaction model). Nest identity was included as a random effect. In the current analysis we have no data on nest volumes but based on field observations the studied *F. aquilonia* and hybrid nests have similar size above ground.

## Results

### Wood ant species maintain their genomic distinctiveness despite hybridisation

We found support for four non-admixed *F. rufa* group species (*F. aquilonia*, *F. lugubris*, *F. rufa* and *F. pratensis*) in our Finland-wide sample based on ADMIXTURE, PCA, and the phylogenetic network (detailed below) (Fig 1, Fig 2, for PCA see Suppl Fig 2). All Finnish *F. polyctena* samples were admixed. Similarly to the genomic data, the morphological analysis identified all studied species. The comparison of ADMIXTURE and NUMOBAT LDA results for non-admixed individuals showed good congruence of genomic data and morphological features (LDA probabilities for non-admixed *F. aquilonia* (N=39) mean=0.93, SD=0.20; *F. polyctena* (N=2) mean=1.00, SD=0.00; *F. rufa* (N=6) mean=0.99, SD=0.02; *F. lugubris* (N=8) mean=1.00, SD=0.00; *F. pratensis* (N=7) mean=0.77, SD=0.30) (Suppl Table 3).

**Figure 2.**
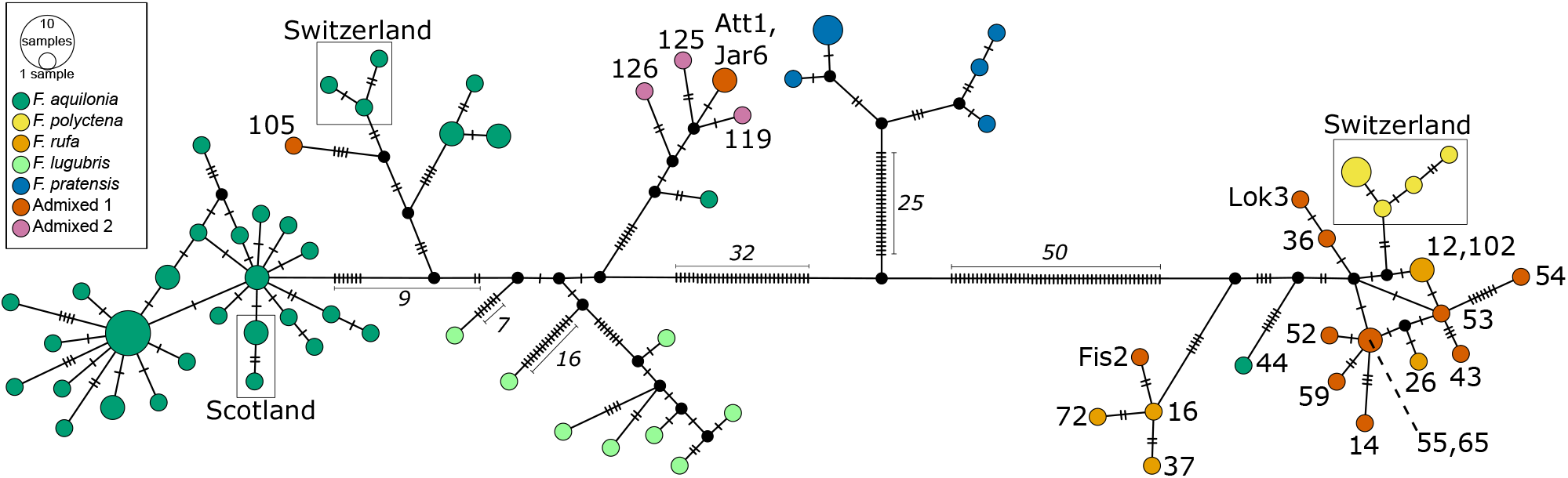
A mitochondrial haplotype network based on 248 SNPs. The circles represent individual haplotypes. The colours indicate nuclear genomic ancestry. The circle sizes indicate the number of identical haplotypes sampled. The dashes represent the number of mutations between the haplotypes, and italicised numbers help illustrate species differences. Pink circles (“Admixed 1”) = admixed individuals between *F. aquilonia* and *F. polyctena/ F. rufa* clades, red circles (“Admixed 2”) = admixed *F. lugubris* individuals. For clarification, selected individual IDs are given. The mitotypes cluster by species, although the difference between *F. polyctena* and *F. rufa* is relatively small given their distant sampling locations. Eleven out of 14 admixed individuals between *F. aquilonia* and the clade of *F. polyctena* and *F. rufa* have mitochondria from the latter southern species.

*Formica pratensis* was genomically most distinct from all the others and formed a well-separated genepool in both nuclear and mitochondrial analyses (Fig 1A&C, Fig 2, Suppl Fig 2A). None of the seven *F. pratensis* populations showed signs of admixture. Further, eight non-admixed *F. lugubris* populations were identified across Finland (Fig 1A&C, Fig 2), their mitochondria similarly clustered together*. F. aquilonia* also separated well from all other species in terms of both nuclear and mitochondrial DNA. Altogether 45 of the *F. aquilonia* populations were found to be non-admixed and to occur across Finland, with relatively low differentiation between mitotypes, considering the great number of samples and wide sampling area. Six non-admixed *F. rufa* populations formed a cluster next to the sister species *F. polyctena* sampled from Switzerland (Fig1 A&C, Fig 2). *F. polyctena,* in contrast, is rare in Finland: no non-admixed *F. polyctena* populations were found from Finland in our study. Since we do not have *F. polyctena* and *F. rufa* from the same location (all *F. polyctena* samples included here are from Switzerland) we cannot exclude the hypothesis that when sympatric they do not form distinct gene pools.

### Extensive hybridisation between *F. aquilonia and the clade of F. polyctena* and *F. rufa, as well as admixed F. lugubris populations*

Our study reveals extensive hybridisation and admixture between sympatric *F. aquilonia*, and the clade of *F. rufa* and *F. polyctena.* The admixed samples always include contribution from the northern species, *F. aquilonia*, and from one or both southern species, *F. polyctena* and/or *F. rufa* (Fig 1A). However, disentangling the relative contributions of *F. polyctena and F. rufa* to the hybrids appears challenging, likely due to their relatively small genomic divergence. The contribution of *F. polyctena* and *F. rufa* to the hybrids vary depending on which non-admixed *F. aquilonia* individuals are chosen for the ADMIXTURE analysis. Admixed populations between *F. aquilonia*, *F. polyctena* and/or *F. rufa* are located in southern Finland, up to a latitude of 62.6°N (Fig 1B). Moreover, all but three (“Att”, “Jar”, “105”) out of these 14 admixed populations have their mitochondria from the southern species. In addition, one individual (“44”) at 65.5°N latitude with a *F. aquilonia* -like nuclear genome showed a mitotype of a southern species (*F. polyctena/ F. rufa clade*), suggesting admixture. Furthermore, three *F. lugubris* populations have indications of admixture with likely contributions from *F. aquilonia*, *F. pratensis* and the *F. polyctena/ F. rufa* clade. One of these hybrid populations is located in southern Finland around latitude 60.2°N, and two in eastern Finland around latitude 62.7°N. In summary, southern Finland represents a mosaic hybrid zone between multiple species, where parental populations are interspersed with hybrid populations.

There was no significant correlation between ADMIXTURE and morphological LDA values in admixed individuals, showing that the relationship of the genotype and phenotype was not linear (Pearson’s correlation of *F. aquilonia* ancestry and respective LDA probability in “Admixed 1” group: r(10) = .36, p = .250; correlation of *F. lugubris* ancestry and respective LDA probability in “Admixed 2” group: r(1) = .96, p = .175).

### Hybrids occur at sites with warmer hibernation temperatures and less spring precipitation than *F. aquilonia* populations

Our results show that the hybrid populations between *F. aquilonia* and the *F. polyctena*/ *F. rufa* clade (“Admixed 1”) occur in locations with warmer temperatures during the ant hibernation period (December-February). The effects of temperature correlated with hybrid occurrence more strongly when warmer extremes in winter (upper quartile, number of non-frost days) were used instead of the average temperature (Fig 3). These results were consistent over measurement periods of 1, 5 and 20 years (see also Suppl Fig 3). Spring precipitation had a strong negative effect on hybrid occurrences over the 20-year period (but not the 1- and 5-year periods), suggesting that hybrid nest locations receive less precipitation in the spring than *F. aquilonia* locations. Latitude and longitude had slight negative effects, reflecting fewer hybrid nest occurrences in the north and the east of the sympatric area than in the south and the west. Neither spring temperature nor winter precipitation had any consistent effects on the probability of hybrids occurrence with our dataset. The estimated effects were weaker, although still significant, for all measurements when the three Åland island nests were removed. The absolute values of these effect estimates are highly correlated with the difference in the expected log pointwise predictive density (elpd-diff) calculated with loo, which was used to rank the models according to their predictive performance (Suppl Fig 4).

**Figure 3.**
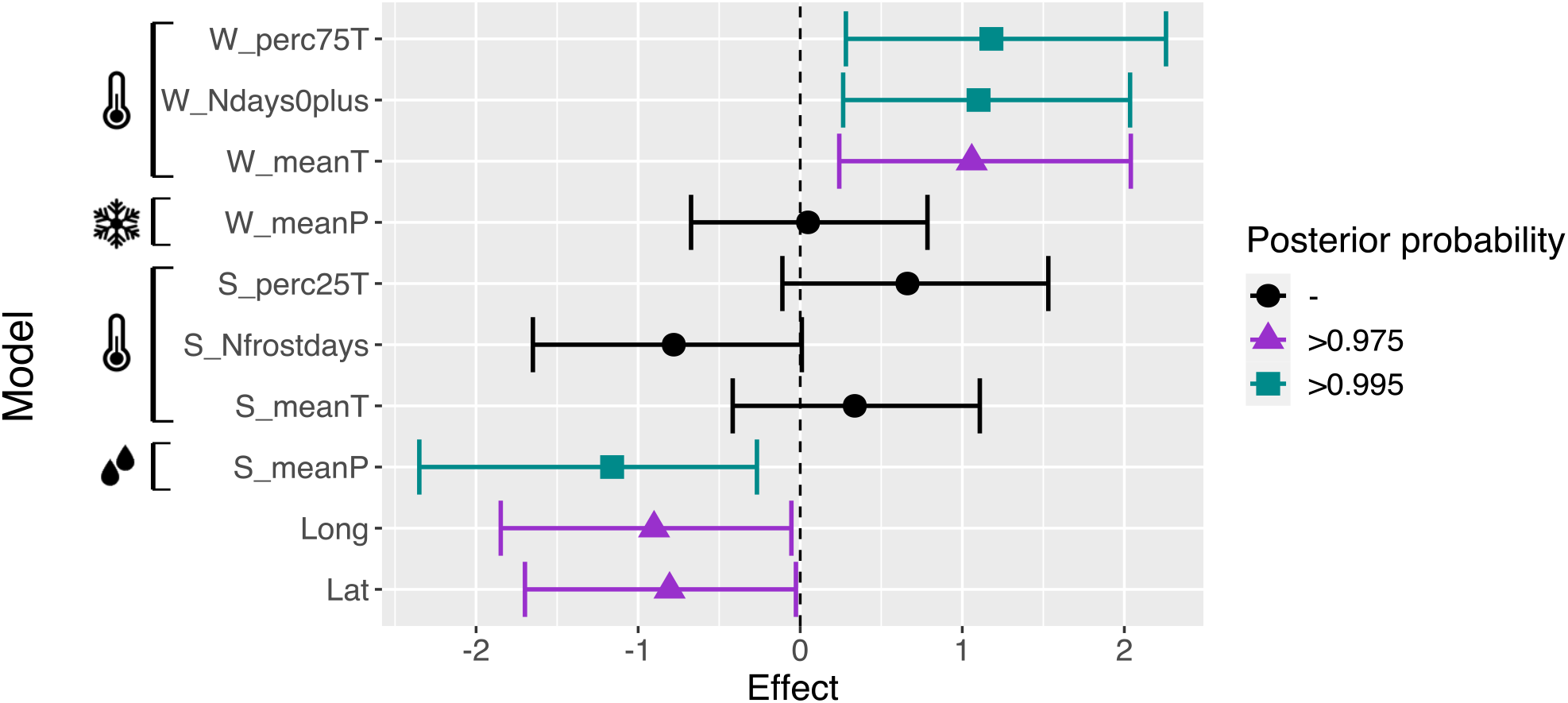
Bayesian logistic regression models show that sympatric *F. aquilonia* and hybrid populations occur in different microclimates, with hybrid nest locations being warmer in the winter and receiving less precipitation in the spring than *F. aquilonia* nest locations. N(nests)=33. Covariates that significantly correlate with hybrid occurrence have non-overlapping credibility intervals (95% credibility intervals are given) with the midline (zero). The winter hibernation season (“W”) = Dec-Feb; spring reproductive season (“S”) = Apr-May. W_perc75T = upper quantile of winter temperatures (T), W_Ndays0plus = number of T > 0°C winter days, W_meanT = annual mean winter T, W_meanP = annual mean winter precipitation, S_perc25T = lower spring T quartile, S_Nfrostdays = number of spring frost days, S_meanT = annual mean spring T, S_meanP = annual mean spring precipitation, Long = longitude, Lat = latitude.

### Ambient temperature impacts within-nest temperature and hybrid nests are warmer than *F. aquilonia* nests

Monitoring of two separate hybrid and two *F. aquilonia* populations shows that ambient temperature has a linear effect on within-nest temperatures in most months (Table 3, Suppl Table 4).

**Table 3.** Interaction model: Bayesian linear mixed model results summary for studying hybrid and *F. aquilonia* within-nest and ambient temperature relationships from two separate study sites, each combining data from the years 2020 and 2021. Absolute effects (in degrees Celsius) and their 95% credibility intervals (CI) are given. * (in boldface) means that the 95 % CI of the estimated effect size does not include 0.

|  | Intercept | Y2021 | Hybrid | ExtT | Hybrid:ExtT | sd(nest) |
| --- | --- | --- | --- | --- | --- | --- |
| Jan | 3.64* (3.14–4.13) | -1.59* (-1.84–(-1.34)) | 0.83* (0.22–1.48) | 0.01 (-0.01–0.04) | 0.08* (0.05–0.12) | 0.97* (0.76–1.25) |
| Feb | 2.95* (2.25–3.64) | -3.37* (-3.71–(-3.03)) | 1.13* (0.23–2.05) | 0.04* (0.01–0.06) | 0.03 (-0.01–0.07) | 1.32* (1.04–1.69) |
| March | 2.90* (1.98–3.85) | -3.05* (-3.38–(-2.71)) | 1.64* (0.40–2.83) | 0.05 (-0.00–0.10) | 0.05 (-0.02–0.12) | 1.94* (1.54–2.45) |
| Apr | 4.40* (2.84–6.02) | -2.39* (-3.06–(-1.74)) | 0.84 (-1.16–2.78) | 0.47* (0.32–0.61) | 0.41* (0.22–0.59) | 3.21* (2.55–4.02) |
| May | 6.52* (4.42–8.62) | -1.93* (-2.62–(-1.24)) | 1.80 (-0.51–4.13) | 0.79* (0.70–0.87) | 0.20* (0.09–0.32) | 4.49* (3.66–5.50) |
| June | 7.83* (5.94–9.87) | -0.22 (-0.56–0.15) | 6.65* (4.30–8.89) | 0.84* (0.79–0.90) | -0.35* (-0.42–(-0.29)) | 3.80* (3.04–4.75) |
| July | 14.67* (12.76–16.55) | -3.17* (-3.60–(-2.75)) | -0.03 (-2.29–2.20) | 0.52* (0.46–0.59) | 0.06 (-0.02–0.15) | 3.51* (2.86–4.34) |

ExtT = Ambient temperature, Y2021 = Year 2021 (2020 is the reference), Hybrid = Nest is hybrid (*F. aquilonia* is the reference), ExtT:Hybrid = Interaction between hybrid and ExtT, sd(nest) = Standard deviation of nest random effect.

Furthermore, even when controlling for the ambient temperature and inter-year effects, hybrid nests were internally warmer, on average, than *F. aquilonia* nests. This result is consistent in both models used (Table 3, Suppl Table 4) from January to March and June. In addition, during January and April-May, hybrid nests warmed up more efficiently with respect to increase in ambient temperatures (see interaction model, Table 3). In June, the hybrid nests tended to warm up more slowly than the cooler *F. aquilonia* nests (Table 3).

### Hybrids show less inbreeding in comparison to parental species

Individuals admixed between *F. aquilonia*, *F. polyctena* and/or *F.rufa* had significantly smaller inbreeding coefficients in comparison to both non-admixed *F. aquilonia* (Wilcoxon Rank Sum test, z=- 4.9, p<0.001) and *F. rufa* (z=-3.38, p<0.001) individuals (Fig 4). This result suggests that hybrids may have a higher adaptive potential.

**Figure 4.**
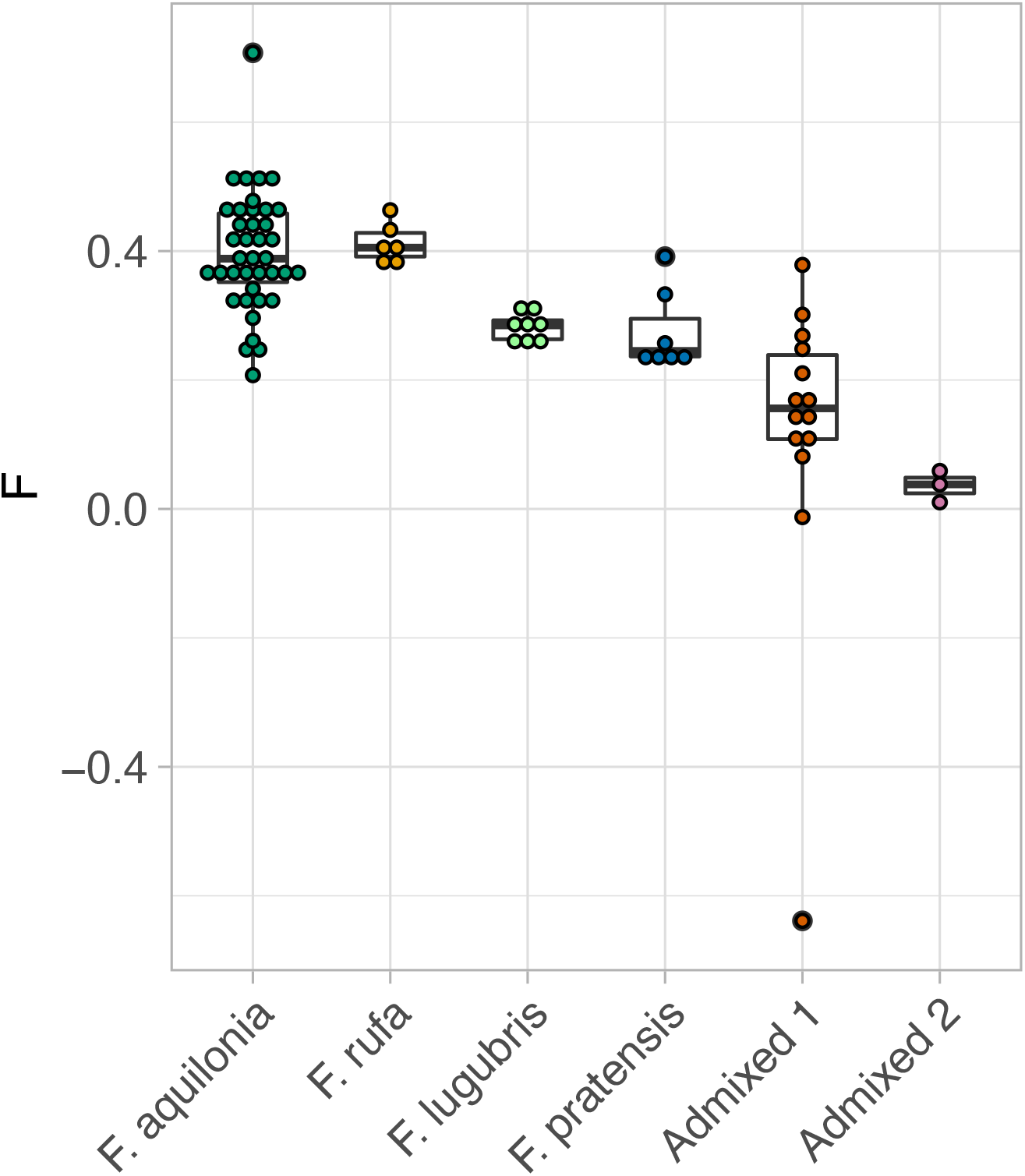
Individual inbreeding coefficients (F) for all Finnish samples show less inbreeding for admixed individuals in comparison to non-admixed samples. Each dot indicates an individual. “Admixed 1” consists of admixed individuals between *F. aquilonia* and *F. polyctena*/ *F. rufa* clade. “Admixed 2” are admixed *F. lugubris* individuals. Inbreeding coefficient medians, first and third quartiles (hinges), and largest and smallest values no further than 1.5 * inter-quartile range from the hinges (whiskers) are shown.

## Discussion

Hybridisation in natural populations is widespread and increased by human activities and a warming climate (Chunco, 2014; Mallet, 2005; Rieseberg, 2009; Scheffers et al., 2016). Consequently, there is a topical need for knowledge on the extent and consequences of hybridisation in natural populations. We report the first genomic study of five *F. rufa* group wood ant species and reveal geographically extensive hybridisation between these diverged species within Finland. We find that all studied members of the *F. rufa* group hybridise in Finland. Three of them form a mosaic hybrid zone in southern Finland with smaller inbreeding coefficients and thus more heterozygosity in hybrids in contrast to parental species. Within this zone admixed populations between *F. aquilonia, F. polyctena* and *F. rufa* are common and interspersed with non-admixed *F. aquilonia* and *F. rufa* populations. All Finnish *F. polyctena* populations analysed in this study are admixed. Within the mosaic zone, the admixed populations occur in habitats that have a higher overwintering temperature and less spring precipitation in comparison to cold-adapted *F. aquilonia*. We further show that hybrid within-nest temperatures are warmer than those of *F. aquilonia* suggesting hybrids and *F. aquilonia* occupy different microclimatic niches.

Extensive hybridisation detected here is likely to impact the ecology and evolution of wood ants. Moreover, hybridisation may allow the different ecological characteristics (Stockan & Robinson, 2016) of the wood ant species to be combined in novel ways in the hybrids. Our results, supported by previous work on thermal responses of the hybrids (Martin-Roy et al., 2021) and on the disadvantages of warm winters for *F. aquilonia* (Sorvari et al., 2011), indicate that rising winter temperatures may favour wood ant hybrids over *F. aquilonia*. In general, our results add important information on the extent of hybridisation in natural populations, and how these populations may cope in changing environments through hybridisation. Our study illustrates how mosaic hybrid zones can have important evolutionary consequences with both ecological and intrinsic selection impacting spatially dispersed independent hybrid populations.

### Genomic distinctness in sympatry, and mosaic hybrid zone with broad implications between differentially adapted *F. rufa* group species

We detected both whole-genome and mitochondrial divergence between species of the *F. rufa* group, as well as hybridisation among all of them in Finland. However, only four out of the five studied species - *F. aquilonia*, *F. lugubris*, *F. rufa* and *F. pratensis* - are found in non-admixed populations in Finland. The mosaic hybrid zone that we detected (ca 60-63 degrees N) greatly expands the known area and extent of wood ant hybridisation in Finland (Beresford et al., 2017; Pamilo & Kulmuni, 2022; Seifert, 2021). We report ten new populations that are admixed between *F. aquilonia* and the *F. polyctena*/ *F. rufa* clade, bringing the total within this dataset to 14. The zone is likely not restricted to our study area, but may continue across the species’ sympatric Eurasian area, as morphological evidence suggests (Seifert, 2021).

The implications and outcomes of mosaic hybrid zones may differ from those of clinal zones. This arises from three features of mosaic zones: i) their patchy nature, ii) the balance of environmentally-dependent and -independent selection and iii) the potential for independent evolution of the hybrid populations when dispersal is limited. All these apply in the *F. rufa* group. First, the differential thermal adaptations (Martin-Roy et al., 2021) and north-south distributions (Stockan & Robinson, 2016) of the parental species are reflected in the patchy distribution of the hybrids and the locally abundant parent. That is, the occurrences of hybrids and parental species are not explained only by a geographic cline as in a clinal hybrid zone model (Arnold, 1997; Barton & Hewitt, 1985), but also by local conditions within each patch within the broad hybrid zone as expected according to the mosaic hybrid zone model (Arnold, 1997).

Second, in contrast to clinal zones, the mosaic zones typically comprise a mixture of environmentally-dependent and -independent selection pressures, the strength of which can vary among patches (Arnold, 1997). The clinal hybrid zones, in contrast, are usually assumed to be arrayed along a linear habitat (Arnold, 1997). The association we observed between hybrids and warmer microclimatic conditions aligns with earlier evidence of thermally-dependent selection (Martin-Roy et al., 2021) that may favour hybrids. This could counteract hybrid breakdown due to intrinsic incompatibilities (Kulmuni et al., 2010; Kulmuni & Pamilo, 2014) and reduced hatching rate (Beresford, 2021) detected previously in wood ant hybrids (Kulmuni et al., 2010; Kulmuni & Pamilo, 2014). Third, the variation in ancestry proportions of *F. aquilonia,* and poor dispersal abilities of supercolonial wood ants, suggest the existence of independent hybrid populations, which has been demonstrated for three closely located populations (Nouhaud et al., 2022). Mosaic of ancestry proportions across the zone contrasts with the spatial gradients typical of clinal hybrid zones (Barton & Hewitt, 1985) and attest that the hybrid populations are long-lived, and thus are not under overwhelmingly negative selection. In summary, evolutionary outcomes of mosaic hybrid zones may differ from those of clinal zones. This is because hybrid populations may evolve independently and be shaped in various ways by both ecological and intrinsic selection pressures and interactions with the parental species leading to a diversity of hybrid lineages and phenotypes (Harrison, 1986; Rand & Harrison, 1989).

Which selection pressures drive the genomic and species composition of the mosaic zone is an interesting question for future studies. They may be, e.g., affected by the degree of forest fragmentation, since in Central Europe *F. polyctena* and *F. rufa* hybrids are more prevalent in small fragments than large forests (Seifert et al., 2010). Furthermore, the interplay of ecological factors such as temperature, habitat preferences, differences in mating times (Martin-Roy et al., 2021; Seifert, 2018; Stockan & Robinson, 2016), and intrinsic selection on genomic incompatibilities (Kulmuni et al., 2010; Kulmuni & Pamilo, 2014) makes predicting the evolutionary dynamics of such a mosaic challenging. Our unique setting of more than ten independent hybrid populations will provide an unrivalled opportunity to study the dynamics and outcomes of hybridization in natural populations.

We predict that the northern border of the hybrid zone, i.e., the northern range limit of *F. rufa* (and *F. polyctena)*, is likely to shift and extend northwards in future following a warming climate, as it has been observed in other taxa (Antão et al., 2022). Global warming is markedly faster in Scandinavia than the global average (Masson-Delmotte et al., 2021), yet biodiversity responses may be complex and not necessarily follow directly biodiversity responses (Waldock et al., 2018).

Our findings are among the few where the mosaic hybrid zone is formed among multiple species. We add a new system to the groups of taxa exhibiting natural mosaic hybrid zones (Abbott & Brennan, 2014; Fraïsse et al., 2014; Rieseberg et al., 1999) and multi-species hybridisation (Banerjee et al., 2022; Grant & Grant, 2020; Heliconius Genome Consortium, 2012; Natola et al., 2022; Ottenburghs, 2019; Reutimann et al., 2020).

### Hybridisation patterns are consistent with the mating strategies of the different species

While hybridisation has been characterised previously in wood ants (Nouhaud et al., 2022; Purcell et al., 2016; Seifert, 2021; Seifert et al., 2010), our study is the first one to report genomic evidence of three-way hybridisation. The parental species have been reported to hybridise only in pairs of *F. aquilonia* × *F. polyctena* and *F. polyctena* × *F. rufa*. Moreover, we show hybridisation between *F. lugubris, F. aquilonia* and other *F. rufa* group species based on three admixed populations from southern and eastern Finland.

The majority of the mosaic zone hybrid populations are of the *F. polyctena*/*F. rufa* clade mitochondrial type, which suggests that hybridisation has mainly occurred between *F. polyctena* or *F. rufa* females and *F. aquilonia* males. This pattern could be caused by life history characteristics of the species and/or due to selection. The simplest explanation is that *F. polyctena* or *F. rufa* queens are forced to mate with *F. aquilonia* males due to lack of conspecifics, because of differences in abundance in Finland (Seifert, 2018), similarly as has been suggested with swordtail fish three-way hybridisation (Banerjee et al., 2022). A complementary explanation is that since highly supercolonial *F. aquilonia* mates on top of or close to its nest (Seifert, 2018), workers protect the queens from non-conspecifics. However, *F. aquilonia* males could disperse further as reported for polygynous *F. lugubris* (Gyllenstrand & Seppä, 2003), which would explain the existence of hybrids. On the contrary, monogyne *F. rufa* queens mate further away from the nests, are not guarded by the workers, and thus may mate more easily with non-conspecifics. Alternatively, the mitochondria of warm-adapted species may be advantageous in environments occupied by hybrid populations (including possibly outcompeting *F. aquilonia*) or mitonuclear incompatibilities may occur when mitochondria comes from *F. aquilonia* but nuclear genome from *F. polyctena* or *F. rufa*. These possibilities remain to be addressed in future studies. The directionality of introgression will be an interesting study question, as throughout the speciation history unidirectional gene flow from *F. aquilonia* to *F. polyctena* has been inferred previously (Portinha et al., 2022).

Our interpretation of clear genomic differentiation between *F. rufa* and *F. polyctena* should be treated with some caution, as all our non-admixed *F. polyctena* populations are from Switzerland. Therefore, we cannot exclude the possibility that sister species *F. rufa* and *F. polyctena* have such a weak RI and substantial gene flow that they cluster genomically together when in sympatry, or resemble different ecotypes of the same species.

### Genomic data is essential in detecting admixture and reveals the rarity of non-admixed *F. polyctena* in Finland

Despite previous morphological evidence, the lack of non-admixed *F. polyctena* within our Finnish samples highlights the importance of integrating genetic data when studying admixture in species that are challenging to distinguish morphologically (Rellstab et al., 2016; Seifert, 2009). The original Finnish National Forest Inventory morphological investigations suggested the presence of *F. polyctena* within Finnish samples (Punttila & Kilpeläinen, 2009b) as did other morphology-based studies (Härkönen & Sorvari, 2014; Sorvari, 2006, 2022; Vepsäläinen & Pisarski, 1982), but it is possible that non-admixed *F. polyctena* does not exist in Finland.

We show that the NUMOBAT morphological analysis can accurately discriminate non-admixed wood ants at the species level. However, in hybrids, the genotype-phenotype relationship is not linear. We demonstrate that a reasonable identification of hybrids and their ancestries by means of NUMOBAT is not possible in multi-species hybridisation. Morphological indication has proven to be a good tool if there is generally a small frequency of hybridization in the material, if the parental species are sufficiently different and if we have a true F1 sample (and not different stages of backcrossing involved). Given these preconditions, good examples for morphological identification of hybrids in ant worker samples are those of *Camponotus herculeanus* × *C. ligniperda* (Seifert, 2019a), *Lasius emarginatus* × *L. platythorax* (Seifert, 2019b) and *Myrmica scabrinodis* × *M. vandeli* (Bagherian Yazdi et al., 2012) or, in case of gyne samples, *Lasius umbratus* × *L. merdionalis* (Seifert, 2006).

The non-linearity of genotype and phenotype can also arise from transgressive segregation: admixed individuals do not necessarily show intermediate phenotypes, but can even exceed phenotypic values of the non-admixed populations due to, e.g., overdominance, epistasis, or complementary action of genes with additive effects (Benowicz et al., 2020). This transgressive segregation has been demonstrated in a variety of taxa including conifers (Benowicz et al., 2020) and flowering plants (Rieseberg et al., 2003; Pisano et al., 2019), but has also been detected in multiple hymenopterans (Gadau et al., 2019; Linksvayer et al., 2009).

### Hybrids occur in warmer and drier habitats and have elevated heterozygosity compared to *F. aquilonia* suggesting they have opportunities to adapt to warming climate

We show that within the mosaic hybrid zone the admixed populations live in habitats with higher overwintering temperatures than those of cold-adapted *F. aquilonia*. Further, we show that ambient and within-nest temperatures are correlated in wood ants, and that hybrid nests are internally warmer than *F. aquilonia* nests.

We cannot rule out non-adaptive explanations for the warmer hybrid habitats, arising e.g. from parental distribution and dispersal patterns. However, the association between hybrids and warmer habitats in our results can arise via selection if *F. aquilonia* suffers more from warm winters due to increased body fat use and a higher metabolic rate. This could lead to decreased survival (Sorvari et al., 2011), or hamper the ants heating the nests in early spring (Seifert, 2018). Thus, hybrid populations could outcompete *F. aquilonia* in warm habitats. In addition, cold winters may reduce hybrid survival, since the studied populations are located at the northern range limits of both *F. polyctena* and *F. rufa*. In a laboratory experiment, *F. aquilonia × F. polyctena* hybrids are less cold-tolerant than non-admixed *F. aquilonia* (Martin-Roy et al., 2021). Both these alternatives align with the hypothesis of better adaptation to warm temperatures in hybrids than in *F. aquilonia.* Although our comparison of ambient spring temperatures between *F. aquilonia* and hybrid habitats across their sympatric zone did not reveal differences, the adaptation hypothesis is supported by hybrid nests being internally warmer and warming up more quickly than *F. aquilonia* nests in the spring. This is when new males and queens are produced by the queen(s), and as such is a crucial time window for the long-term maintenance of the colony. Wood ants can actively control their within-nest temperatures, and during springtime the within-nest temperature deviates most significantly from the ambient temperature (Frouz & Finer, 2007). This may make ambient spring temperatures less crucial for ant survival than winter temperatures. These results suggest that ambient temperature fluctuations might affect the wood ants’ hibernation and reproductive success. It will be of interest in future studies to determine whether and to what extent thermal differences detected between *F. aquilonia* and hybrid nests are caused by either differences in the worker population size, ambient temperature, relative sun exposure, nest structure, and/or ant activity patterns.

The longer-term adaptive potential of the hybrids in a changing environment is further supported by our finding of less inbreeding in hybrids compared to the non-admixed individuals, as genetic diversity could provide means to adapt to new environmental challenges (Grant & Grant, 2019; Seehausen, 2013).

## Conclusions

We used combined genomic and morphological approaches to analyse species divergence and hybridisation in natural populations of extensively hybridising (Seifert, 2021), evolutionarily young (Goropashnaya et al., 2004, 2012) boreal keystone species. This first genomic study on these five *F. rufa* group wood ant species shows that they form their own gene pools, yet we cannot exclude the possibility that *F. rufa* and *F. polyctena* are two different ecotypes rather than species. We further reveal a mosaic hybrid zone in southern Finland, composed of three interbreeding wood ant species with differing climatic preferences. Our results align with other recent studies proposing that hybridisation is more prevalent in natural populations than previously thought. Moreover, they support a hypothesis that the wood ant hybrids may harbour adaptive potential (Martin-Roy et al., 2021) in the face of an increasingly warming northern climate (Masson-Delmotte et al., 2021), which affects species’ distributions and requires adaptation to new conditions (Antão et al., 2022). This is because we show that the hybrids live in warmer environments than the sympatric non-admixed *F. aquilonia* populations. Since the majority of our samples were collected over 15 years ago, repeated sampling of the same and novel locations could provide insights into the evolution of the mosaic hybrid zone and pinpoint the role of climate as a potential driver of its evolution. Future research is needed to examine the relative roles and patterns of barriers to gene flow between the wood ant species, as well as patterns of introgression, and consequences of hybridisation for the wood ants and the surrounding ecosystem.

## Acknowledgements

We thank Natural Resources Institute Finland for access to samples collected during the 10th National Forest Inventory (NFI) of Finland, and Leena Finér and Jouni Kilpeläinen for organising the collection of the NFI ant data. Timo Domisch, Pirjo Appelgren, Gergely Várkonyi, Riitta Savolainen and Kari Vepsäläinen provided additional ant samples. We would also like to thank Raphael Martin-Roy for helping to collect the within-nest temperature data, Jaakko Kuurne for involvement in the preliminary mitochondrial data analyses, and the SpecIAnt research group for comments and support. Warm thanks for Martyn James and other Biomedicum Functional Genomics Unit personnel for expertise and advice in genome sequencing, and CSC–ITCenter for Science, Finland, for computational resources. Mirkka Jones and Jukka Sirén acknowledge support from the Academy of Finland's ‘Thriving Nature’ research profiling action.

## Data availability

SNP and climate data will be made available via Dryad and Figshare upon publication.

## Author contributions

Original idea - IS, JK

Conceptualisation - IS, PP, HH, JK.

Methodology and Investigation - IS, PN, BS, RS, MJ, JS.

Funding acquisition - IS, JK.

Project administration - IS, JK.

Study material and resources - BS, PP, JK.

Supervision - HH, JK.

Visualisation - IS.

Writing - original draft - IS, MJ, JS, HH, JK.

Writing - review & editing - IS, PN, PP, BS, RS, MJ, JS, HH, JK.

All authors approved the manuscript before submission.

## References

Aalto, J., Pirinen, P., & Jylhä, K. (2016). New gridded daily climatology of Finland: Permutation-based uncertainty estimates and temporal trends in climate. Journal of Geophysical Research, D: Atmospheres, 121(8). https://doi.org/10.1002/2015JD024651

Abbott, R. J., & Brennan, A. C. (2014). Altitudinal gradients, plant hybrid zones and evolutionary novelty. In Philosophical Transactions of the Royal Society B: Biological Sciences (Vol. 369, Issue 1648, p. 20130346). https://doi.org/10.1098/rstb.2013.0346

Alexander, D. H., & Lange, K. (2011). Enhancements to the ADMIXTURE algorithm for individual ancestry estimation. BMC Bioinformatics, 12, 246.

Alexander, D. H., Novembre, J., & Lange, K. (2009). Fast model-based estimation of ancestry in unrelated individuals. Genome Research, 19(9), 1655–1664.

Ålund, M., Immler, S., Rice, A. M., & Qvarnström, A. (2013). Low fertility of wild hybrid male flycatchers despite recent divergence. Biology Letters, 9(3), 20130169.

Antão, L. H., Weigel, B., Strona, G., Hällfors, M., Kaarlejärvi, E., Dallas, T., Opedal, Ø. H., Heliölä, J., Henttonen, H., Huitu, O., Korpimäki, E., Kuussaari, M., Lehikoinen, A., Leinonen, R., Lindén, A., Merilä, P., Pietiäinen, H., Pöyry, J., Salemaa, M., … Laine, A.-L. (2022). Climate change reshuffles northern species within their niches. Nature Climate Change, 12(6), 587–592.

Arnold, M. L. (1997). Natural Hybridization and Evolution (Oxford Series in Ecology and Evolution) (1st ed.). Oxford University Press.

Bagherian Yazdi, A., Münch, W., & Seifert, B. (2012). A first demonstration of interspecific hybridization in Myrmica ants by geometric morphometrics (Hymenoptera: Formicidae). Myrmecological News / Osterreichische Gesellschaft Fur Entomofaunistik, 17, 121–131.

Banerjee, S. M., Powell, D. L., Moran, B. M., Ramírez-Duarte, W. F., Langdon, Q. K., Gunn, T. R., Vazquez, G., Rochman, C., & Schumer, M. (2022). Complex hybridization between deeply diverged fish species in a disturbed ecosystem. In bioRxiv (p. 2022.10.08.511445). https://doi.org/10.1101/2022.10.08.511445

Barton, N. H., & Hewitt, G. M. (1985). Analysis of Hybrid Zones. Annual Review of Ecology and Systematics, 16(1), 113–148.

Benowicz, A., Stoehr, M., Hamann, A., & Yanchuk, A. D. (2020). Estimation of the F2 generation segregation variance and relationships among growth, frost damage, and bud break in coastal Douglas-fir (Pseudotsuga menziesii (Mirb.) Franco) wide-crosses. Annals of Forest Science, 77(2), 1–13.

Beresford, J. (2021). The role of hybrids in the process of speciation : a study of naturally occurring Formica wood ant hybrids [Helsingin yliopisto]. https://helda.helsinki.fi/handle/10138/335389

Beresford, J., Elias, M., Pluckrose, L., Sundström, L., Butlin, R. K., Pamilo, P., & Kulmuni, J. (2017). Widespread hybridization within mound-building wood ants in Southern Finland results in cytonuclear mismatches and potential for sex-specific hybrid breakdown. Molecular Ecology, 26(15), 4013–4026.

Bolger, A. M., Lohse, M., & Usadel, B. (2014). Trimmomatic: a flexible trimmer for Illumina sequence data. Bioinformatics, 30(15), 2114–2120.

Brelsford, A., Purcell, J., Avril, A., Tran Van, P., Zhang, J., Brütsch, T., Sundström, L., Helanterä, H., & Chapuisat, M. (2020). An Ancient and Eroded Social Supergene Is Widespread across Formica Ants. Current Biology: CB, 30(2), 304–311.e4.

Bryant, D., & Moulton, V. (2004). Neighbor-net: an agglomerative method for the construction of phylogenetic networks. Molecular Biology and Evolution, 21(2), 255–265.

Bürkner, P.-C. (2017). brms: An R Package for Bayesian Multilevel Models Using Stan. Journal of Statistical Software, 80, 1–28.

Masson-Delmotte, V., P. Zhai, A. Pirani, S.L. Connors, C. Péan, S. Berger, N. Caud, Y. Chen, L. Goldfarb, M.I. Gomis, M. Huang, K. Leitzell, E. Lonnoy, J.B.R. Matthews, T.K. Maycock, T. Waterfield, O. Yelekçi, R. Yu, and B. Zhou (eds.). (2021). IPCC, 2021: Climate Change 2021: The physical science basis. Contribution of Working Group I to the Sixth Assessment Report of the Intergovernmental Panel on Climate Change. Cambridge University Press, Cambridge, United Kingdom and New York, NY, USA, 2391 pp. doi:10.1017/9781009157896.

Chunco, A. J. (2014). Hybridization in a warmer world. Ecology and Evolution, 4(10), 2019–2031.

Danecek, P., Auton, A., Abecasis, G., Albers, C. A., Banks, E., DePristo, M. A., Handsaker, R. E., Lunter, G., Marth, G. T., Sherry, S. T., McVean, G., Durbin, R., & 1000 Genomes Project Analysis Group. (2011). The variant call format and VCFtools. Bioinformatics, 27(15), 2156–2158.

Danecek, P., Bonfield, J. K., Liddle, J., Marshall, J., Ohan, V., Pollard, M. O., Whitwham, A., Keane, T., McCarthy, S. A., Davies, R. M., & Li, H. (2021). Twelve years of SAMtools and BCFtools. GigaScience, 10(2). https://doi.org/10.1093/gigascience/giab008

De-Kayne, R., Selz, O. M., Marques, D. A., Frei, D., Seehausen, O., & Feulner, P. G. D. (2022). Genomic architecture of adaptive radiation and hybridization in Alpine whitefish. Nature Communications, 13(1), 4479.

Ellison, C. K., Niehuis, O., & Gadau, J. (2008). Hybrid breakdown and mitochondrial dysfunction in hybrids of Nasonia parasitoid wasps. Journal of Evolutionary Biology, 21(6), 1844–1851.

Fraïsse, C., Roux, C., Welch, J. J., & Bierne, N. (2014). Gene-flow in a mosaic hybrid zone: is local introgression adaptive? Genetics, 197(3), 939–951.

Frouz, J. (2000). The effect of nest moisture on daily temperature regime in the nests of Formica polyctena wood ants. Insectes Sociaux, 47(3), 229–235.

Frouz, J., & Finer, L. (2007). Diurnal and seasonal fluctuations in wood ant (Formica polyctena) nest temperature in two geographically distant populations along a south – north gradient. Insectes Sociaux, 54(3), 251–259.

Gadau, J., Pietsch, C., Gerritsma, S., Ferber, S., van de Zande, L., van den Assem, J., & Beukeboom, L. W. (2019). Genetic architecture of male courtship behavior differences in the parasitoid wasp genus Nasonia (Hymenoptera; Pteromalidae). In bioRxiv (p. 831735). https://doi.org/10.1101/831735

Garrison, E., Kronenberg, Z. N., Dawson, E. T., Pedersen, B. S., & Prins, P. (2022). A spectrum of free software tools for processing the VCF variant call format: vcflib, bio-vcf, cyvcf2, hts-nim and slivar. PLoS Computational Biology, 18(5), e1009123.

Garrison, E., & Marth, G. (2012). Haplotype-based variant detection from short-read sequencing. In arXiv [q-bio.GN]. arXiv. http://arxiv.org/abs/1207.3907

Goropashnaya, A. V., Fedorov, V. B., & Pamilo, P. (2004). Recent speciation in the Formica rufa group ants (Hymenoptera, Formicidae): inference from mitochondrial DNA phylogeny. Molecular Phylogenetics and Evolution, 32(1), 198–206.

Goropashnaya, A. V., Fedorov, V. B., Seifert, B., & Pamilo, P. (2012). Phylogenetic relationships of Palaearctic Formica species (Hymenoptera, Formicidae) based on mitochondrial cytochrome B sequences. PloS One, 7(7), e41697.

Grant, P. R., & Grant, B. R. (2019). Hybridization increases population variation during adaptive radiation. Proceedings of the National Academy of Sciences of the United States of America, 116(46), 23216–23224.

Grant, P. R., & Grant, B. R. (2020). Triad hybridization via a conduit species. Proceedings of the National Academy of Sciences of the United States of America, 117(14), 7888–7896.

Gyllenstrand, N., & Seppä, P. (2003). Conservation genetics of the wood ant, Formica lugubris, in a fragmented landscape. Molecular Ecology, 12(11), 2931–2940.

Härkönen, S. K., & Sorvari, J. (2014). Species richness of associates of ants in the nests of red wood antFormica polyctena(Hymenoptera, Formicidae). Insect Conservation and Diversity / Royal Entomological Society of London, 7(6), 485–495.

Harrison, R. G. (1986). Pattern and process in a narrow hybrid zone. Heredity, 56(3), 337–349.

Heliconius Genome Consortium. (2012). Butterfly genome reveals promiscuous exchange of mimicry adaptations among species. Nature, 487(7405), 94–98.

Horstmann, K., & Schmid, H. (1986). Temperature regulation in nests of the wood ant, Formica polyctena (Hymenoptera: Formicidae). Entomologia Generalis, 11(3-4), 229–236.

Huson, D. H., & Bryant, D. (2006). Application of phylogenetic networks in evolutionary studies. Molecular Biology and Evolution, 23(2), 254–267.

Kadochová, Š., & Frouz, J. (2014). Red wood ants Formica polyctena switch off active thermoregulation of the nest in autumn. Insectes Sociaux, 61(3), 297–306.

Kagawa, K., & Takimoto, G. (2018). Hybridization can promote adaptive radiation by means of transgressive segregation. Ecology Letters, 21(2), 264–274.

Katoh, K., & Standley, D. M. (2013). MAFFT multiple sequence alignment software version 7: improvements in performance and usability. Molecular Biology and Evolution, 30(4), 772–780.

Kulmuni, J., & Pamilo, P. (2014). Introgression in hybrid ants is favored in females but selected against in males. Proceedings of the National Academy of Sciences of the United States of America, 111(35), 12805–12810.

Kulmuni, J., Seifert, B., & Pamilo, P. (2010). Segregation distortion causes large-scale differences between male and female genomes in hybrid ants. Proceedings of the National Academy of Sciences of the United States of America, 107(16), 7371–7376.

Leigh, J. W., & Bryant, D. (2015). Popart: Full-feature software for haplotype network construction. Methods in Ecology and Evolution / British Ecological Society, 6(9), 1110–1116.

Lemmon, E. M., & Juenger, T. E. (2017). Geographic variation in hybridization across a reinforcement contact zone of chorus frogs (Pseudacris). Ecology and Evolution, 7(22), 9485–9502.

Leroy, T., Louvet, J.-M., Lalanne, C., Le Provost, G., Labadie, K., Aury, J.-M., Delzon, S., Plomion, C., & Kremer, A. (2020). Adaptive introgression as a driver of local adaptation to climate in European white oaks. The New Phytologist, 226(4), 1171–1182.

Li, H. (2013). Aligning sequence reads, clone sequences and assembly contigs with BWA-MEM. In arXiv [q-bio.GN]. arXiv. http://arxiv.org/abs/1303.3997

Linksvayer, T. A., Rueppell, O., Siegel, A., Kaftanoglu, O., Page, R. E., Jr, & Amdam, G. V. (2009). The genetic basis of transgressive ovary size in honeybee workers. Genetics, 183(2), 693–707, 1SI – 13SI.

Mallet, J. (2005). Hybridization as an invasion of the genome. Trends in Ecology & Evolution, 20(5), 229–237.

Martin-Roy, R., Nygård, E., Nouhaud, P., & Kulmuni, J. (2021). Differences in Thermal Tolerance between Parental Species Could Fuel Thermal Adaptation in Hybrid Wood Ants. The American Naturalist, 198(2), 278–294.

Meier, J. I., Marques, D. A., Mwaiko, S., Wagner, C. E., Excoffier, L., & Seehausen, O. (2017). Ancient hybridization fuels rapid cichlid fish adaptive radiations. Nature Communications, 8, 14363.

Natola, L., Seneviratne, S. S., & Irwin, D. (2022). Population genomics of an emergent tri-species hybrid zone. In bioRxiv (p. 2022.06.04.494703). https://doi.org/10.1101/2022.06.04.494703

Nouhaud, P., Beresford, J., & Kulmuni, J. (2021). Cost-effective long-read assembly of a hybrid *Formica aquilonia* × *Formica polyctena* wood ant genome from a single haploid individual. In bioRxiv. bioRxiv. https://doi.org/10.1101/2021.03.09.434597

Nouhaud, P., Martin, S. H., Portinha, B., Sousa, V. C., & Kulmuni, J. (2022). Rapid and repeatable genome evolution across three hybrid ant populations. In bioRxiv (p. 2022.01.16.476493). https://doi.org/10.1101/2022.01.16.476493

Ottenburghs, J. (2019). Multispecies hybridization in birds. Avian Research, 10(1), 1–11.

Owens, G. L., & Samuk, K. (2019). Adaptive introgression during environmental change can weaken reproductive isolation. Nature Climate Change, 10(1), 58–62.

Pamilo, P., & Kulmuni, J. (2022). Genetic identification of Formica rufa group species and their putative hybrids in northern Europe. Myrmecological News / Osterreichische Gesellschaft Fur Entomofaunistik, 32. https://www.biotaxa.org/mn/article/view/76072

Pedersen, B. S., & Quinlan, A. R. (2018). Mosdepth: quick coverage calculation for genomes and exomes. Bioinformatics, 34(5), 867–868.

Pfennig, K. S. (2016). Reinforcement as an initiator of population divergence and speciation. Current Zoology, 62(2), 145–154.

Portinha, B., Avril, A., Bernasconi, C., Helanterä, H., Monaghan, J., Seifert, B., Sousa, V. C., Kulmuni, J., & Nouhaud, P. (2022). Whole-genome analysis of multiple wood ant population pairs supports similar speciation histories, but different degrees of gene flow, across their European ranges. Molecular Ecology, 31(12), 3416–3431.

Punttila, P., & Kilpeläinen, J. (2009). Distribution of mound-building ant species (Formica spp., Hymenoptera) in Finland: preliminary results of a national survey. Annales Zoologici Fennici, 46(1), 1–15.

Purcell, J., Zahnd, S., Athanasiades, A., Türler, R., Chapuisat, M., & Brelsford, A. (2016). Ants exhibit asymmetric hybridization in a mosaic hybrid zone. Molecular Ecology, 25(19), 4866–4874.

Purcell, S., Neale, B., Todd-Brown, K., Thomas, L., Ferreira, M. A. R., Bender, D., Maller, J., Sklar, P., de Bakker, P. I. W., Daly, M. J., & Sham, P. C. (2007). PLINK: a tool set for whole-genome association and population-based linkage analyses. American Journal of Human Genetics, 81(3), 559–575.

Rand, D. M., & Harrison, R. G. (1989). Ecological Genetics of a Mosaic Hybrid Zone: Mitochondrial, Nuclear, and Reproductive Differentiation of Crickets by Soil Type. Evolution; International Journal of Organic Evolution, 43(2), 432–449.

Rellstab, C., Bühler, A., Graf, R., Folly, C., & Gugerli, F. (2016). Using joint multivariate analyses of leaf morphology and molecular-genetic markers for taxon identification in three hybridizing European white oak species (Quercus spp.). Annals of Forest Science, 73(3), 669–679.

Reutimann, O., Gugerli, F., & Rellstab, C. (2020). A species-discriminatory single-nucleotide polymorphism set reveals maintenance of species integrity in hybridizing European white oaks (Quercus spp.) despite high levels of admixture. Annals of Botany, 125(4), 663–676.

Rieseberg, L. H. (2009). Evolution: replacing genes and traits through hybridization. Current Biology: CB, 19(3), R119–R122.

Rieseberg, L. H., Raymond, O., Rosenthal, D. M., Lai, Z., Livingstone, K., Nakazato, T., Durphy, J. L., Schwarzbach, A. E., Donovan, L. A., & Lexer, C. (2003). Major ecological transitions in wild sunflowers facilitated by hybridization. Science, 301(5637), 1211–1216.

Rieseberg, L. H., Whitton, J., & Gardner, K. (1999). Hybrid zones and the genetic architecture of a barrier to gene flow between two sunflower species. Genetics, 152(2), 713–727.

Rubini Pisano, A., Moré, M., Cisternas, M. A., Raguso, R. A., & Benitez-Vieyra, S. (2019). Breakdown of species boundaries in Mandevilla: floral morphological intermediacy, novel fragrances and asymmetric pollen flow. Plant Biology, 21(2), 206–215.

Scheffers, B. R., De Meester, L., Bridge, T. C. L., Hoffmann, A. A., Pandolfi, J. M., Corlett, R. T., Butchart, S. H. M., Pearce-Kelly, P., Kovacs, K. M., Dudgeon, D., Pacifici, M., Rondinini, C., Foden, W. B., Martin, T. G., Mora, C., Bickford, D., & Watson, J. E. M. (2016). The broad footprint of climate change from genes to biomes to people. Science, 354(6313). https://doi.org/10.1126/science.aaf7671

Seehausen, O. (2013). Conditions when hybridization might predispose populations for adaptive radiation [Review of *Conditions when hybridization might predispose populations for adaptive radiation*]. Journal of Evolutionary Biology, 26(2), 279–281.

Seifert, B. (2006). Social cleptogamy in the ant subgenus Chthonolasius--survival as a minority. Abhandlungen Und Berichte Des Naturkundemuseums Görlitz, 77, 251–276.

Seifert, B. (2009). Cryptic species in ants (Hymenoptera: Formicidae) revisited: we need a change in the alpha-taxonomic approach. Myrmecological News / Osterreichische Gesellschaft Fur Entomofaunistik, 12(12), 149–166.

Seifert, B. (2018). The ants of central and north Europe. lutra Verlags-und Vertriebsgesellschaft.

Seifert, B. (2019a). Lasius nigroemarginatus Forel, 1874 is a F1 Hybrid between L. emarginatus (Olivier, 1792) and L. platythorax Seifert, 1991 (Hymenoptera, Formicidae). Beiträge Zur Entomologie= Contributions to Entomology, 69(2), 291–300.

Seifert, B. (2019b). Hybridization in the European carpenter ants Camponotus herculeanus and C. ligniperda (Hymenoptera: Formicidae). Insectes Sociaux, 66(3), 365–374.

Seifert, B. (2021). A taxonomic revision of the Palaearctic members of the Formica rufa group (Hymenoptera: Formicidae) – the famous mound-building red wood ants. Myrmecological News / Osterreichische Gesellschaft Fur Entomofaunistik, 31. https://www.biotaxa.org/mn/article/view/68533

Seifert, B., Kulmuni, J., & Pamilo, P. (2010). Independent hybrid populations of Formica polyctena X rufa wood ants (Hymenoptera: Formicidae) abound under conditions of forest fragmentation. Evolutionary Ecology, 24(5), 1219–1237.

Sorvari, J. (2006). Two distinct morphs in the wood ant *Formica polyctena* in Finland: a result of hybridization? Entomologica Fennica, 17(1), 1–7.

Sorvari, J. (2022). Biogeography and habitat preferences of red wood ants of the Formica rufa group (Hymenoptera: Formicidae) in Finland, based on citizen science data. European Journal of Entomology, 119, 92–98.

Sorvari, J., Haatanen, M.-K., & Vesterlund, S.-R. (2011). Combined effects of overwintering temperature and habitat degradation on the survival of boreal wood ant. Journal of Insect Conservation, 15(5), 727–731.

Stockan, J. A., & Robinson, E. J. H. (2016). Wood Ant Ecology and Conservation. Cambridge University Press.

Tan, A., Abecasis, G. R., & Kang, H. M. (2015). Unified representation of genetic variants. Bioinformatics, 31(13), 2202–2204.

Toyama, K. S., Crochet, P.-A., & Leblois, R. (2020). Sampling schemes and drift can bias admixture proportions inferred by structure. Molecular Ecology Resources, 20(6), 1769–1785.

Trigos-Peral, G., Juhász, O., Kiss, P. J., Módra, G., Tenyér, A., & Maák, I. (2021). Wood ants as biological control of the forest pest beetles Ips spp. Scientific Reports, 11(1), 17931.

Vehtari, A., Gelman, A., & Gabry, J. (2017). Practical Bayesian model evaluation using leave-one-out cross-validation and WAIC. Statistics and Computing, 27(5), 1413–1432.

Vepsäläinen, K., & Pisarski, B. (1982). Assembly of island ant communities. Annales Zoologici Fennici, 19(4), 327–335.

Waldock, C., Dornelas, M., & Bates, A. E. (2018). Temperature-Driven Biodiversity Change: Disentangling Space and Time. Bioscience, 68(11), 873–884.

