## Supplemental Figures for "Extensive hybridisation between multiple differently adapted species may aid persistence in a changing climate"

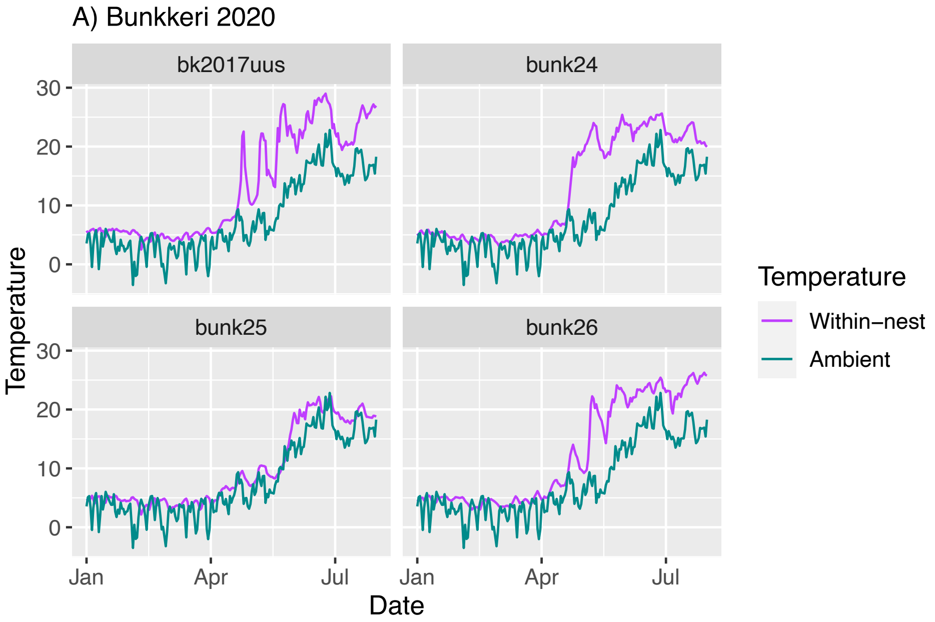


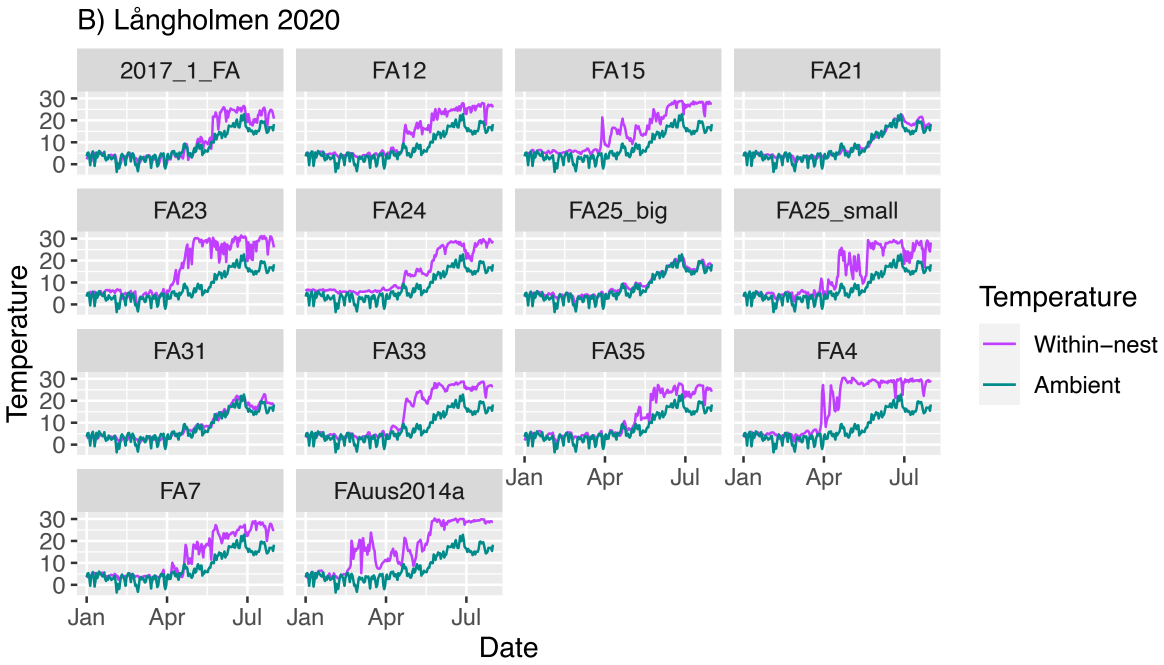

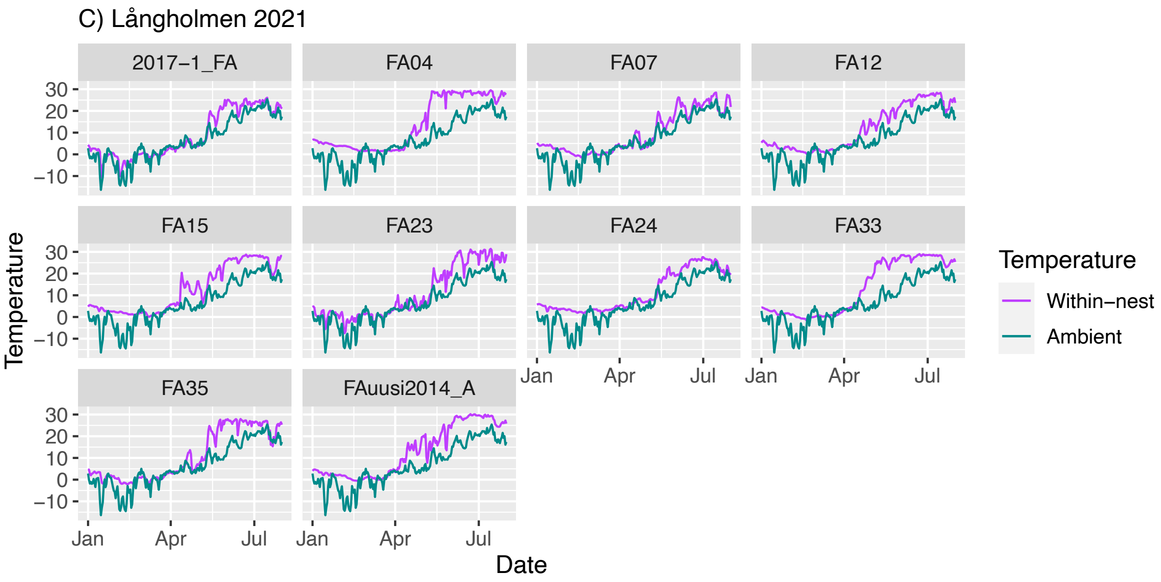
C) Långholmen 2021
2017−1_FA
FA04
FA07
FA12
FA15
FA23
FA24
FA33
FA35
FAuusi2014_A
Temperature
Date
Jan
Apr
Jul
Temperature
Within−nest
Ambient

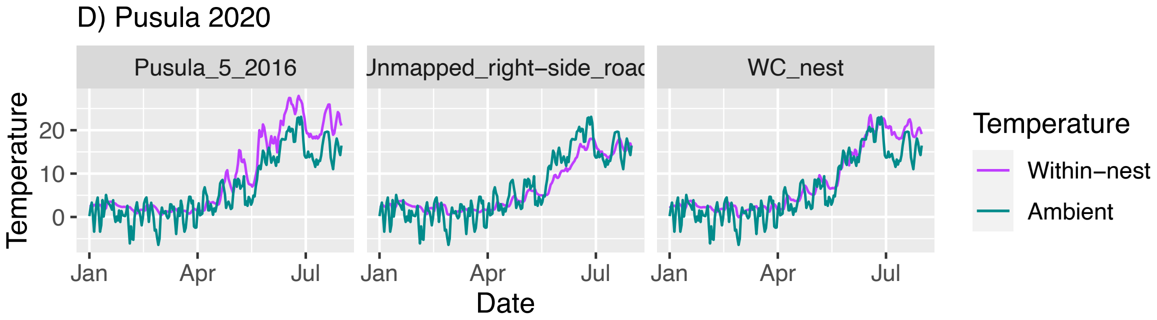
D) Pusula 2020
Pusula_5_2016
Jnmapped_right−side_roac
WC_nest
Temperature
Date
Jan
Apr
Jul
Temperature
Within−nest
Ambient

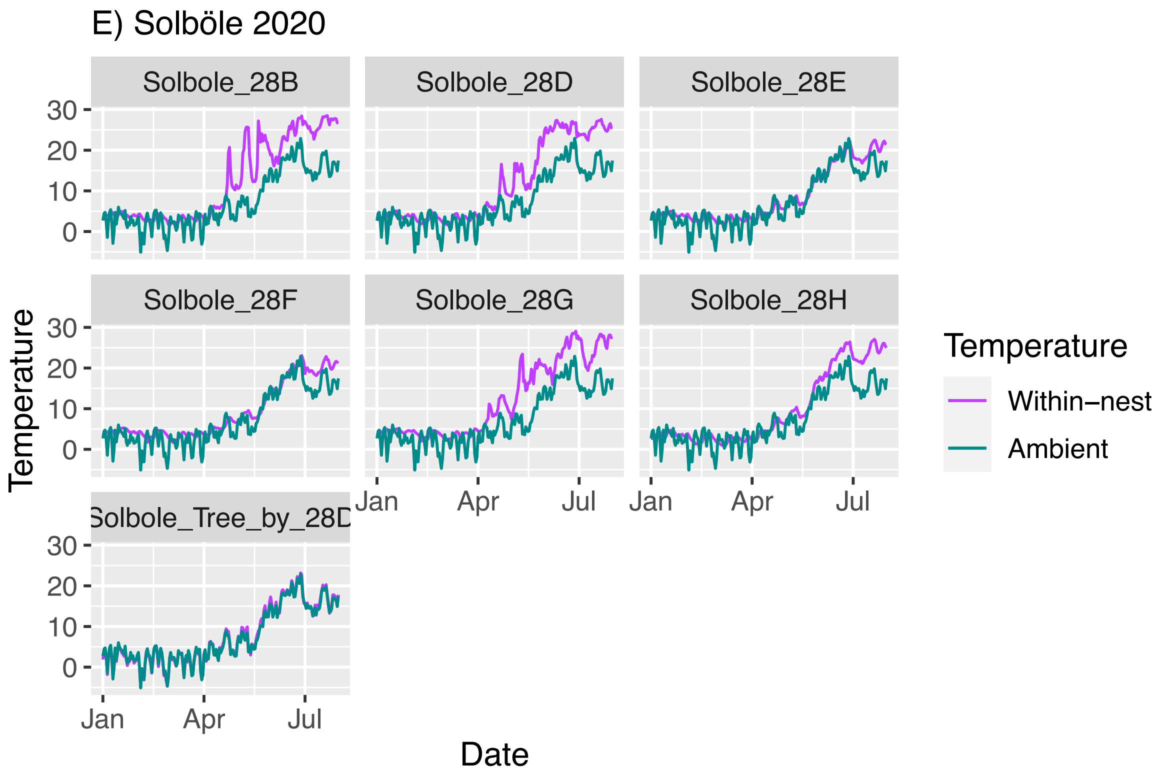
E) Solböle 2020
Solbole_28B
Solbole_28D
Solbole_28E
Solbole_28F
Solbole_28G
Solbole_28H
Solbole_Tree_by_28D
Temperature
Date
Jan
Apr
Jul
Temperature
Within−nest
Ambient

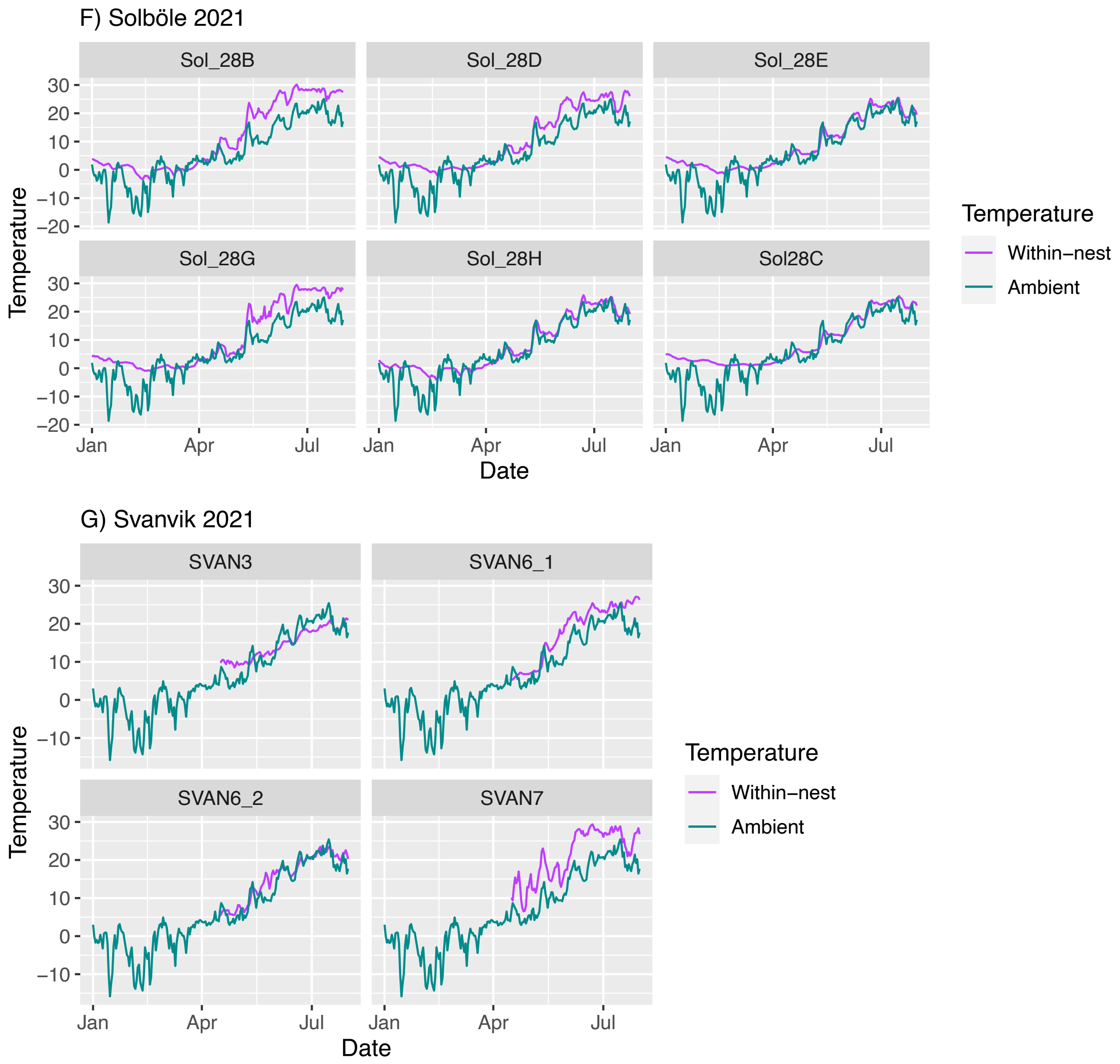


**Supplementary Figure 1:** Absolute within- & ambient temperatures for each studied nest: Bunkkeri hybrid population 2020 (A), Långholmen hybrid population 2020 (B) and 2021 (C), Pusula *F. aquilonia* population 2020 (D) , Solböle *F. aquilonia* population 2020 (E) and 2021 (F), and Svanvik hybrid population 2021 (G).

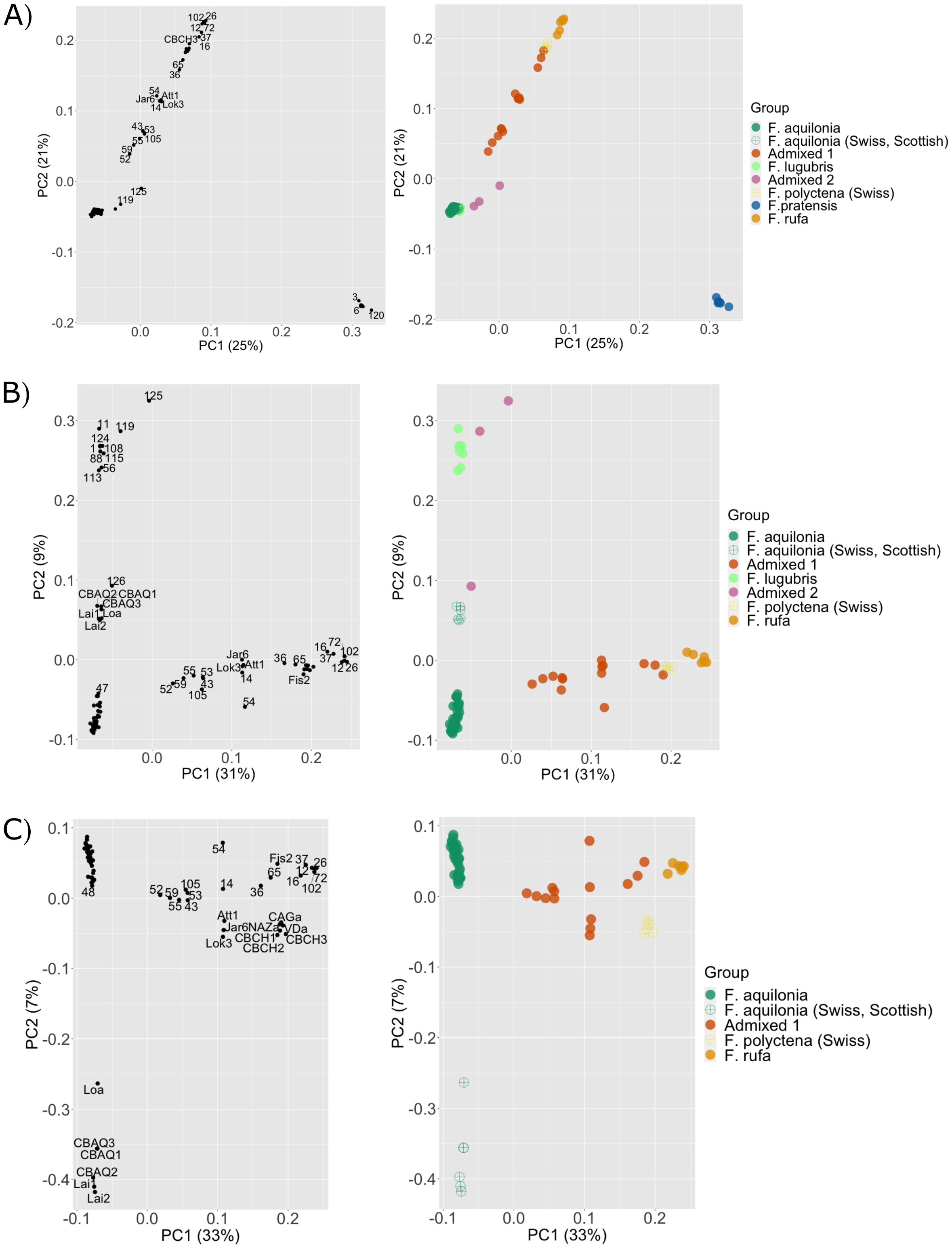


**Supplementary Figure 2:** Principal component analysis (PCA) on all samples (A), and with increased resolution either without *F. pratensis* samples (B) or without both *F. pratensis* and *F. lugubris* samples (and their hybrids) (C). Left side plots provide sample identity information.

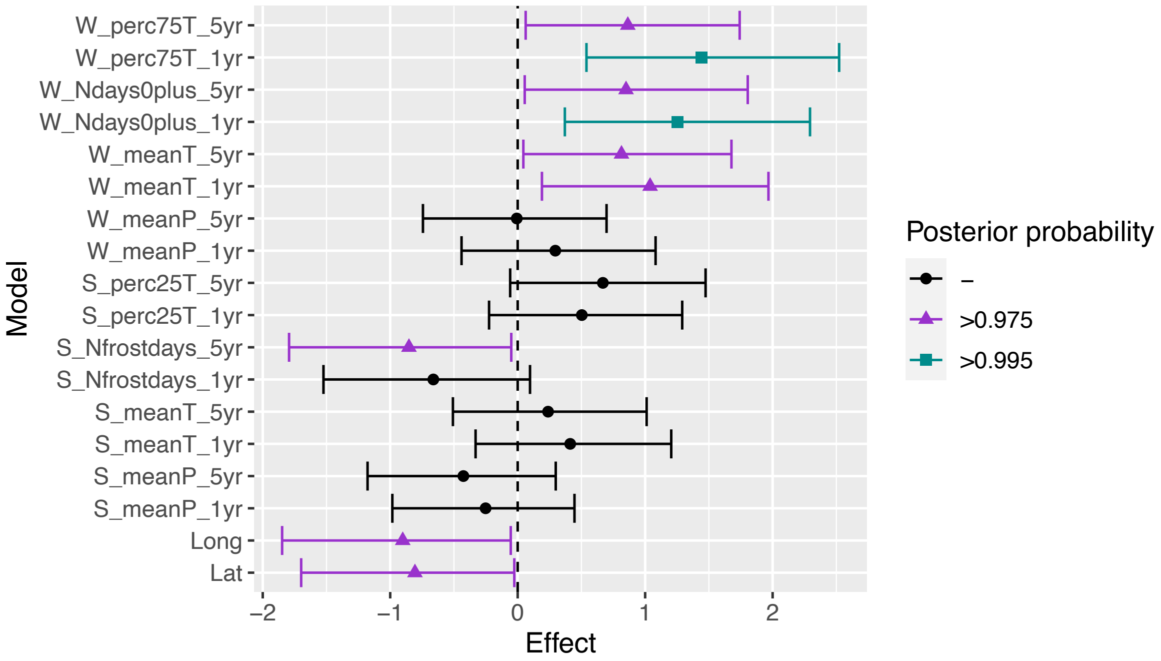


**Supplementary Figure 3:** Bayesian logistic regression climatic analysis results for 1- and 5-year spans. N(nests)=33. Covariates that significantly correlate with hybrid occurrence have non-overlapping credibility intervals (95% credibility intervals are given) with the midline (zero). The winter hibernation season ("W") = Dec-Feb; spring reproductive season ("S") = Apr-May. W_perc75T = upper quantile of winter temperatures (T), W_Ndays0plus = number of T > 0°C winter days, W_meanT = annual mean winter T, W_meanP = annual mean winter precipitation, S_perc25T = lower spring T quartile, S_Nfrostdays = number of spring frost days, S_meanT = annual mean spring T, S_meanP = annual mean spring precipitation, Long = longitude, Lat = latitude.

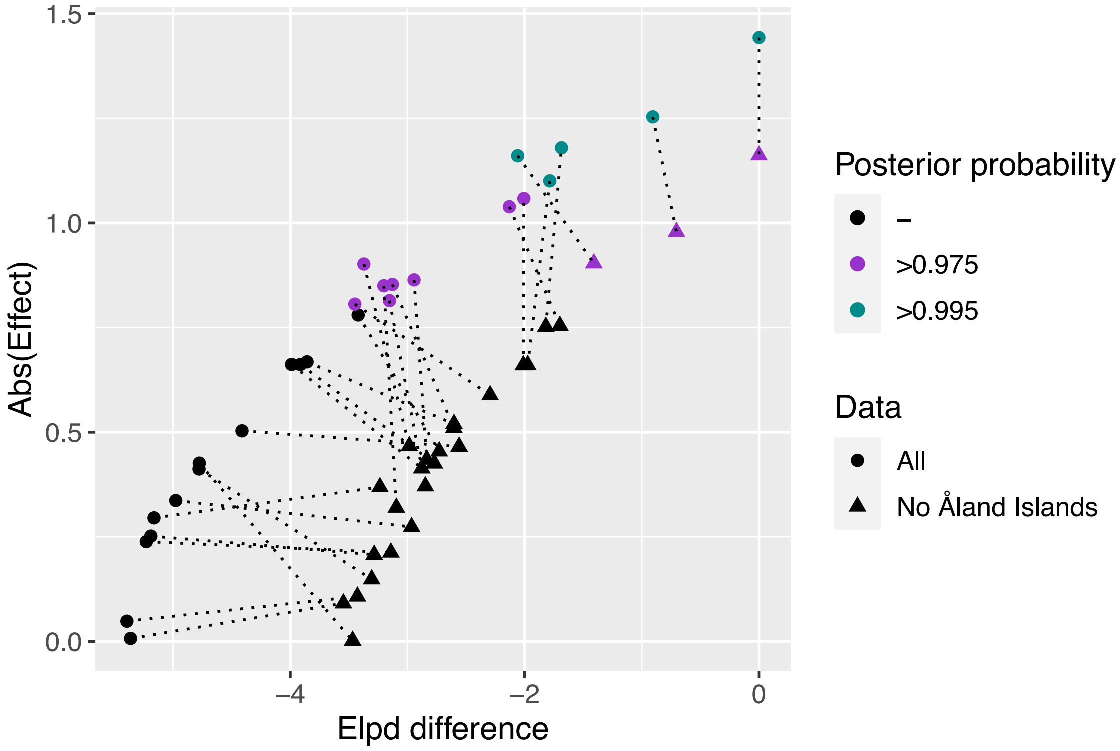


**Supplementary Figure 4:** The difference in the Bayesian logistic regression model expected log pointwise predictive density (elpd-diff) calculated with leave one out (loo) cross-validation is highly correlated with the absolute value of the effect estimates.
